# Mitochondrial dysfunction triggers actin polymerization necessary for rapid glycolytic activation

**DOI:** 10.1101/2022.06.03.494723

**Authors:** Rajarshi Chakrabarti, Tak Shun Fung, Taewook Kang, Pieti W. Pallijeff, Anu Suomalainen, Edward J. Usherwood, Henry N. Higgs

**Author notes:** These authors contributed equally to this work (Rajarshi Chakrabarti, Tak Shun Fung).

## Abstract

Mitochondrial damage represents a dramatic change in cellular homeostasis, necessitating rapid responses. One rapid response is peri-mitochondrial actin polymerization, termed ADA (<u>a</u>cute <u>d</u>amage-induced <u>a</u>ctin). The consequences of ADA are not fully understood. Here we show that ADA is necessary for rapid glycolytic activation upon inhibition of mitochondrial ATP production in multiple cells, including mouse embryonic fibroblasts and effector CD8^+^ T lymphocytes, for which glycolysis is an important source of ATP and biosynthetic molecules. Treatments that induce ADA include CCCP, antimycin A, rotenone, oligomycin, and hypoxia. The Arp2/3 complex inhibitor CK666 or the mitochondrial sodium-calcium exchanger (NCLX) inhibitor CGP37157, applied simultaneously with the ADA stimulus, inhibit both ADA and the glycolytic increase within 5-min, suggesting that ADA is necessary for glycolytic stimulation. Two situations causing chronic reductions in mitochondrial ATP production, ethidium bromide treatment (to deplete mitochondrial DNA) and mutation to the NDUFS4 subunit of complex 1 of the electron transport chain, cause persistent peri-mitochondrial actin filaments of similar morphology to ADA. Both peri-mitochondrial actin loss and a 20% ATP decrease occur within 10 min of CK666 treatment in NDUFS4 knock-out cells. We propose that ADA is necessary for rapid glycolytic activation upon mitochondrial impairment, to re-establish ATP production.

## INTRODUCTION

Mitochondrial damage represents an acute cellular stress, compromising ATP production and the balance of several key metabolites, as well as a rise in reactive oxygen species in some situations(Kasahara and Scorrano, 2014; Nunnari and Suomalainen, 2012). Cells respond in many ways to mitochondrial damage, including up-regulating glycolysis and mitochondrial destruction by mitophagy (Pickles et al., 2018; Sturdik et al., 1986). These responses require extensive communication between mitochondria and the rest of the cell, and defects in these responses are linked to multiple pathologies such as Parkinson’s.

One response is ADA (<u>a</u>cute <u>d</u>amage-induced <u>a</u>ctin), resulting in a dense actin filament network surrounding the mitochondrion (Fung et al., 2019; Li et al., 2015). This actin network is dependent on Arp2/3 complex, and has morphological similarities to other Arp2/3 complex-dependent mitochondrial polymerization events that occur in interphase (Moore et al., 2016) and mitotic cells (Moore et al., 2021). ADA is distinct, however, from another population of actin filaments that influences mitochondria, which we call CIA (calcium-induced actin). CIA is not dependent on Arp2/3 complex, but instead on the formin INF2, which is activated by increased cytoplasmic calcium (Chakrabarti et al., 2018; Ji et al., 2015; Shao et al., 2015; Wales et al., 2016). A consequence of CIA is increased mitochondrial division, through two mechanisms: 1) increased ER-to-mitochondrial calcium transfer, leading to increased inner mitochondrial membrane (IMM) dynamics; and 2) increased recruitment of the mitochondrial division factor Drp1 to the outer mitochondrial membrane (OMM), leading to increased OMM dynamics (Chakrabarti et al., 2018).

The function of ADA is at present unclear. One study suggests that ADA increases mitochondrial division (Li et al., 2015). Our previous data, however, suggest that ADA actually decreases the extensive mitochondrial dynamics that occur in the acute stages of mitochondrial depolarization (Fung et al., 2022; Fung et al., 2019). We also show that these mitochondrial dynamics are more consistent with changes to IMM morphology, driven by the IMM protease Oma1, than they are with mitochondrial division. These findings are in line with several previous studies (De Vos et al., 2005; Liu and Hajnóczky, 2011; Minamikawa et al., 1999; Miyazono et al., 2018). The acute mitochondrial changes induced by depolarization are independent of Drp1 (Fung et al., 2019; Miyazono et al., 2018).

Here, we provide evidence for a second function for ADA: stimulation of the rapid increase in glycolysis that occurs after mitochondrial ATP production is compromised. This effect on glycolysis occurs upon a variety of treatments, including hypoxia. Based on these results, we postulate that ADA represents an acute response to maintain cellular ATP levels in the face of mitochondrial dysfunction.

## RESULTS

To begin our investigation into the function of ADA, we first asked whether ADA is a common cellular response, both in terms of cell type and in terms of the nature of the mitochondrial assault. ADA is rapidly and transiently induced by CCCP, a mitochondrial depolarizer, in multiple cell lines including mouse embryonic fibroblasts (MEFs), U2-OS, HeLa and Cos-7 cells (Fig. 1; and Movies 1-4). In all cases, maximum actin polymerization occurs within 4 min, and actin is largely depolymerized in 10 min. Closer examination shows that actin accumulates around most but not all mitochondria, in both live-cell imaging of multiple cell types (Fig. 1A) and fixed-cell imaging of MEFs (Fig. S1). The actin-free mitochondria are frequently smaller, which may be due to CCCP-induced circularization that has been previously identified (De Vos et al., 2005; Liu and Hajnóczky, 2011; Minamikawa et al., 1999; Miyazono et al., 2018) and that we have shown to be inhibited by ADA (Fung et al., 2022; Fung et al., 2019).

**Figure 1.**
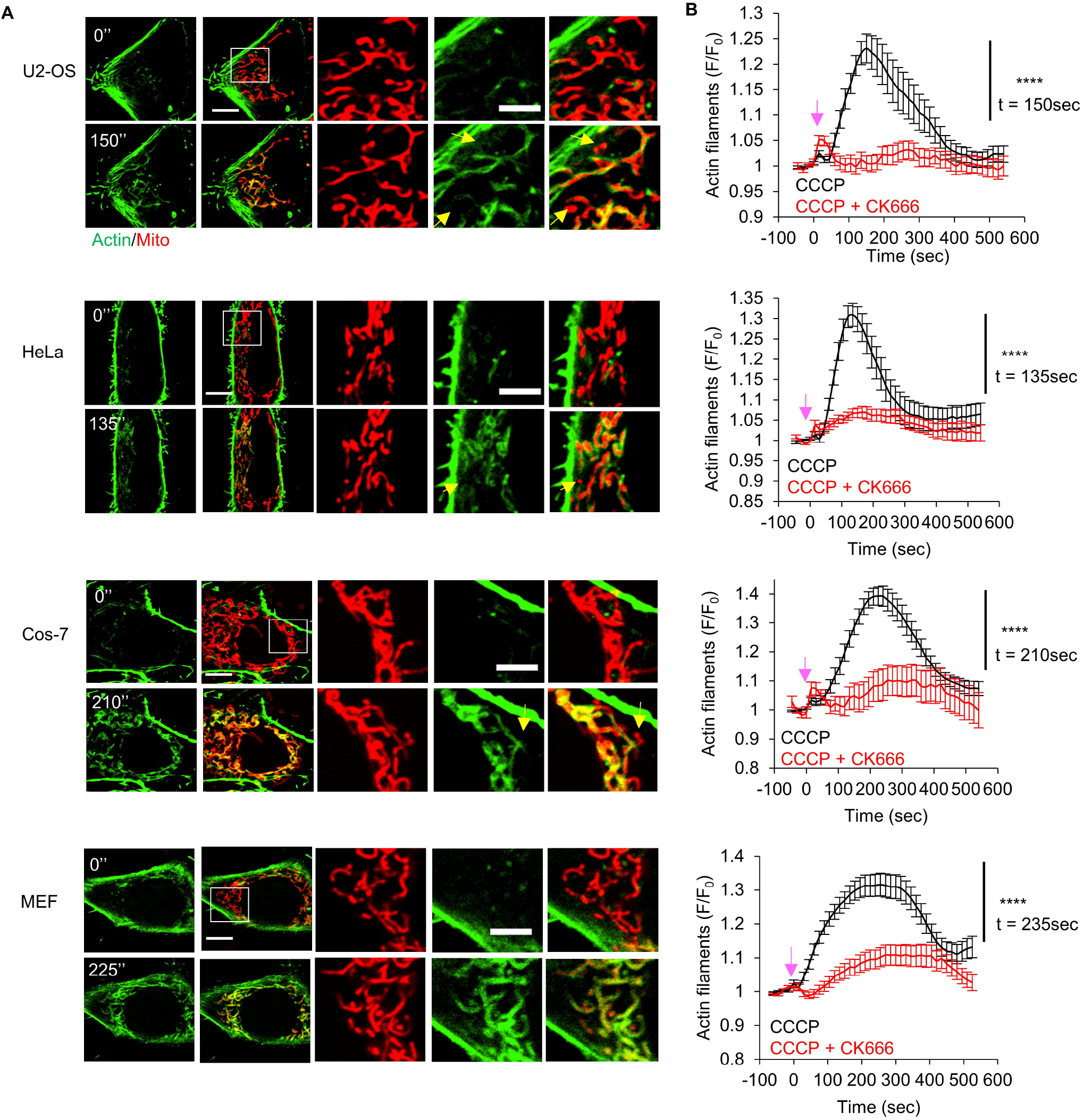
ADA in multiple cell types. A) Micrographs of live-cell imaging for U2-OS, HeLa, Cos-7 and MEF cell before (0 sec) and at their peak ADA timepoints after 20μM CCCP addition (150sec – U2-OS; 135sec - HeLa; 210sec – Cos-7; 225sec – MEF). All cells were transfected with markers for actin filaments (GFP-Ftractin, green) and mitochondria (mito-DsRed, red). Scale bars are 10 μm (full cell), 5 μm (inset). Yellow arrow indicates punctate mitochondrion without actin assembly. Corresponds to movie 1-4. B) Graph of actin intensity (± s.e.m.) around mitochondria in U2-OS, HeLa, Cos7 and MEF cells as a function of time of 20μM CCCP or 100μM CK666 + 20μM CCCP simultaneous treatment. Cells per condition: U2-OS, n = 31; HeLa, n = 32; Cos-7, n ≥ 17; MEF, n ≥ 22. Arrows indicate time of treatment. **** P < 0.0001. Experiments done in 25mM glucose with serum. Exact number of experiments, statistical tests and sample sizes are provided in Supplementary Table 1.

Treatment with CK666, an Arp2/3 complex inhibitor (Nolen et al., 2009), inhibits ADA in all cell types tested (Fig. 1 B), similar to our previous results in U2-OS cells (Fung et al., 2022; Fung et al., 2019) and to the actin “waves” recently shown in interphase and mitotic cells (Moore et al., 2021; Moore et al., 2016). ADA does not appear to drive directional mitochondrial motility (Movies 1-4), and the actin polymerization rarely extends appreciably beyond the mitochondrion (Movie 5), in contrast to the motility-inducing actin ‘tails’ previously shown to assemble from actin clouds in mitotic cells (Moore et al., 2021).

Since CCCP is a relatively harsh treatment, resulting in complete mitochondrial depolarization in seconds, we tested two electron transport chain (ETC) inhibitors: antimycin A (Complex III) and rotenone (Complex I), which cause partial depolarization, as measured by tetramethylrhodamine ethyl ester (TMRE) (Fig. 2 A). ADA is induced within 3 min for both antimycin A and rotenone, in a CK666-inhibitable manner (Fig. 2, B and C). Importantly, CK666 effectively inhibits ADA when added simultaneously to the ADA stimulus, suggesting that the effect of Arp2/3 complex inhibition is on the acute ADA response. We also examined the effect of the ATP synthase inhibitor oligomycin on ADA, which causes a slight increase in mitochondrial polarization over 5 min (Fig. S2 A). Oligomycin stimulates ADA in a CK666-inhibitable manner (Fig. S2, B and C). This result suggests that ADA is not triggered by decreased mitochondrial polarization.

**Figure 2.**
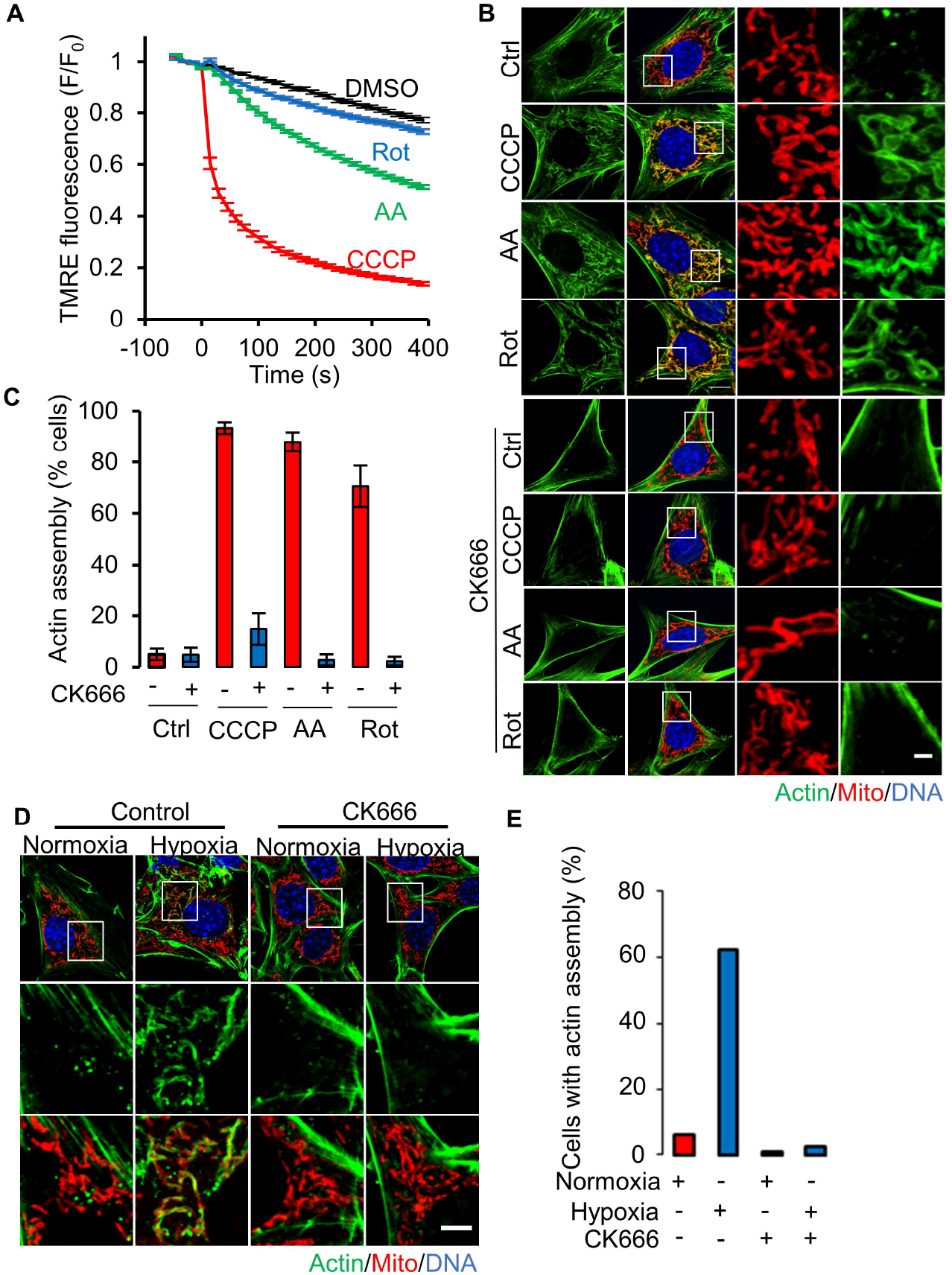
ADA stimulation by mitochondrial depolarization or ETC inhibition. A) Mitochondrial depolarization (assessed by TMRE fluorescence) in MEFs with DMSO, CCCP, antimycin A or rotenone (± s.e.m.) treatments. n ≥ 98 cells per group. Experiments done in 25mM glucose with serum. B) MEFs stained for actin filaments (TRITC-phalloidin, green), mitochondria (Tom20, red) and DNA (DAPI, blue) after 3 min treatment with DMSO, 20μM CCCP, 25μM antimycin A or 50μM rotenone in the absence (top) or presence (bottom) of 100μM CK666. Right images are zooms of boxed regions. Scale bar: 5 μm. C) % cells (± s.d.) displaying ADA for the conditions shown in B. n ≥ 62/18 cells/fields of view (FOV) per group. Experiments done in 25mM glucose with serum. D) MEFs stained similarly to B, in normoxia or hypoxia for 30 min, in presence of absence of 100μM CK-666. Scale bar: 5 μm. E) % cells displaying ADA after 30 min normoxia or hypoxia, in the absence or presence of 100μM CK-666. n ≥ 174/20 cells/FOV per group. Experiments done in 25mM glucose without serum. Exact number of experiments, FOV and sample size are provided in Supplementary Table 1.

Another deleterious treatment is hypoxia, which depletes a necessary substrate for Complex IV of the ETC. Upon exposure to hypoxia (1% oxygen), morphologically similar actin filaments to those generated by the ADA treatments arise within 30 min (Fig. 2, D and E). Hypoxia-induced actin polymerization is inhibited by CK666 (Fig. 2, D and E). These results show that ADA is a rapid response to multiple acute treatments that inhibit oxidative phosphorylation (oxphos), including chemical treatments (CCCP, antimycin A, rotenone, oligomycin) and oxygen deprivation (hypoxia).

### ADA is required for rapid up-regulation of glycolysis upon oxphos inhibition

What might be the function of ADA? Since ADA is stimulated by treatments that inhibit oxphos, we asked whether inhibiting ADA would have an impact on cytoplasmic ATP levels. For these experiments, we used the GO-ATeam1 ATP biosensor (Nakano et al., 2011) in live MEFs. To inhibit ADA, we used CK666 added simultaneously to the stimulus, decreasing the possibility of longer-term CK666 effects. We conducted the experiment at two glucose concentrations: 25 mM, which is the concentration in DMEM but is ∼5-fold higher than serum glucose; and 2 mM, which is hypoglycemic compared to serum but is similar to the extracellular glucose concentration a number of environments, including solid tumors (Ho et al., 2015) and in the brain (Silver and Erecińska, 1994). ADA occurs in MEFs in hypoglycemic conditions (Fig. S2 D), similar to our earlier results in 25 mM glucose (Fig. 1 A, B).

At 2 mM glucose, there is a 20% drop in ATP within 2 min of mitochondrial depolarization by CCCP. Simultaneous addition of CK666 increases the ATP drop to >30% (Fig. 3 A; and Fig. S3 A). Biochemical assays of whole-cell ATP levels show similar results (Fig. S3 B). The effects of antimycin A or rotenone on ATP levels are slower than for CCCP, with the rotenone effect being negligible (Fig. 3 B; and Fig. S3 C). However, CK666 addition causes significant additional drops in cytosolic ATP for both antimycin A and rotenone treatment at 2 mM glucose (Fig. 3 B; and Fig. S3 C). At 25 mM glucose, CK666 has a non-significant effect on ATP levels when added with CCCP, antimycin A or rotenone (Fig. S3, D and E). Treatment with CK666 alone does not have a significant effect on ATP levels at either glucose concentration (Fig. 3, A and B; and Fig. S3, D and E). These experiments suggest that ADA is necessary to maintain cellular ATP levels upon oxphos inhibition when glucose is limited.

**Figure 3.**
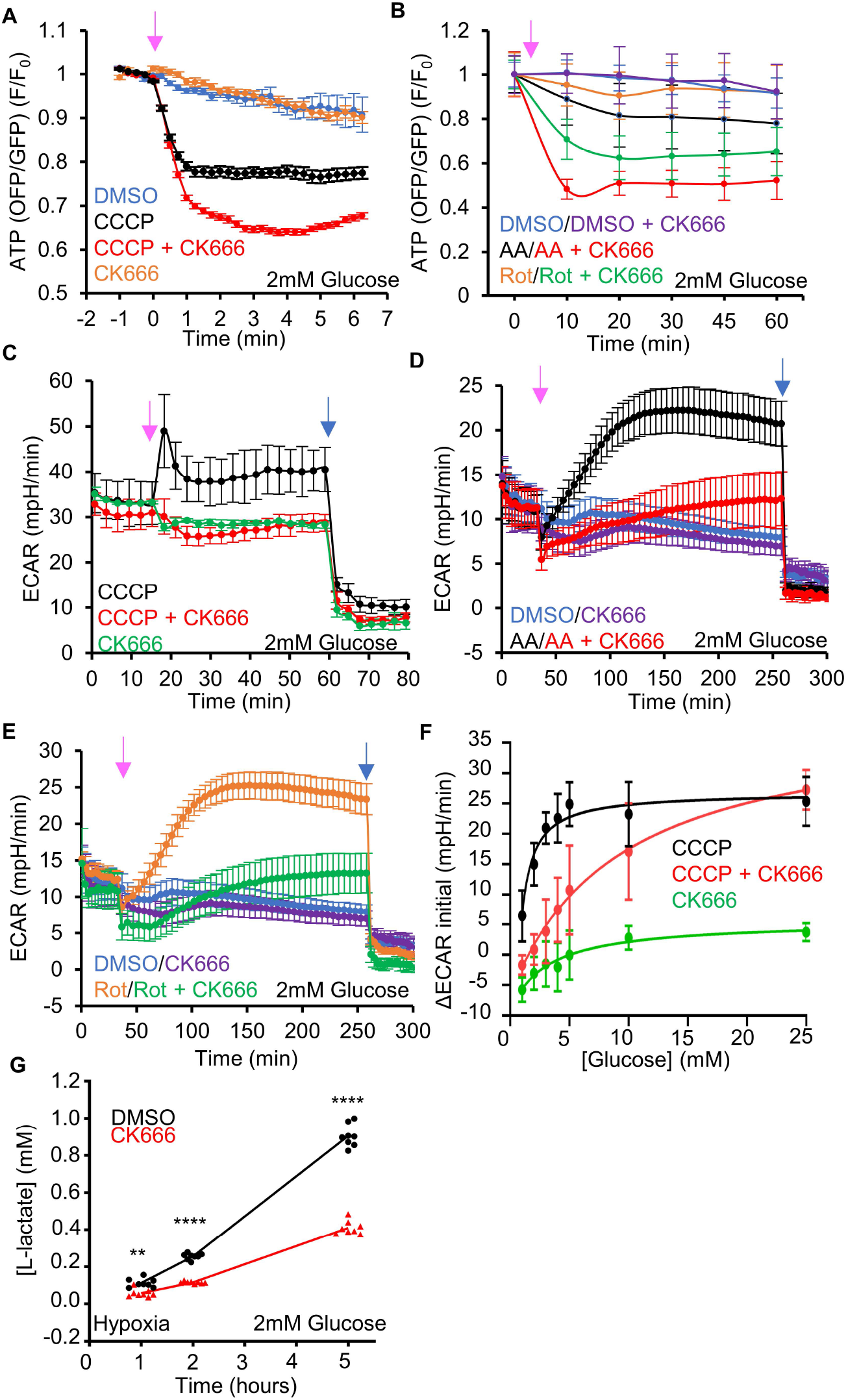
ADA is required for glycolytic activation upon mitochondrial perturbation in MEFs. A) Cytoplasmic ATP levels (± s.e.m.) after 20μM CCCP in the absence or presence of 100μM CK666, using GO-ATeam1. n ≥ 35 cells per group. P values graphed in Fig. S3 A. Arrow indicates time of treatment. Experiments done in 2mM glucose with serum. B) Cytoplasmic ATP levels (± s.e.m.) after 25μM antimycin A or 50μM rotenone in the absence or presence of 100μM CK666, using GO-ATeam1. n ≥ 24 cells per group. P values graphed in Fig. S3 C. Arrow indicates time of treatment. Experiments done in 2mM glucose with serum. C) ECAR (± s.d.) upon 100μM CK666, 1μM CCCP or 1μM CCCP + 100μM CK666 addition (15 min), followed by 50mM 2-deoxyglucose (2-DG) (59 min) in 2mM glucose medium without serum. Pink arrow indicates drug treatment and blue arrow indicate 2-DG treatment. D) ECAR (± s.d.) upon DMSO, 100μM CK666, 2.5μM antimycin A or 2.5μM antimycin A + 100μM CK666 addition (33 min), then 50mM 2-DG (258 min) in 2mM glucose medium without serum. Pink arrow indicates drug treatment and blue arrow indicate 2-DG treatment. E) ECAR (± s.d.) upon DMSO, 100μM CK666, 5μM rotenone or 5μM rotenone + 100μM CK666 addition (33 min), then 50mM 2-DG (258 min) in 2mM glucose medium without serum. Pink arrow indicates drug treatment and blue arrow indicate 2-DG treatment. F) Effect of glucose concentration on ECAR spike (± s.d.) induced by 3 min 1μM CCCP, with and without 100μM CK666. P values graphed in Fig. S4 D. G) Effect of 100μM CK666 on lactate production in hypoxia (1% O_2_) in MEFs at 2mM glucose without serum. Points indicate individual well measurements starting with 100,000 cells/well. ** P = 0.0018. **** P < 0.0001. Number of experiments, FOV, sample sizes and statistical tests are provided in Supplementary Table 1.

Inhibition of oxphos causes an increase in glycolysis to make up for decreased ATP production (Krebs, 1972; Racker, 1974). Changes in glycolysis can be assayed by changes in extracellular acidification rate (ECAR), an indirect measure of lactate production (Mookerjee et al., 2017). Treatment of MEFs with CCCP causes a rapid ECAR spike followed by prolonged ECAR elevation in both 2 mM and 25 mM glucose medium (Fig. 3 C; and Fig. S4 A). The initial ECAR spike occurs at the first measurable timepoint after CCCP addition (3-min). Antimycin A and rotenone also induce ECAR increases, but not as rapidly as CCCP (Fig. 3, D and E; and Fig. S4, B and C).

For all three treatments, addition of CK666 simultaneously with the treatment suppresses the ECAR increase in 2 mM glucose (Fig. 3, C-E) but not in 25 mM glucose (Fig. S4, A-C). Titrating the glucose concentration, we find significant effects of CK666 on ECAR occur at 5 mM glucose and below for CCCP treatment, for both the initial effect (3 min, Fig. 3 F; and Fig. S4 D), or at 40 min after treatment (Fig. S4, E and F). These results show that Arp2/3 complex-mediated actin polymerization is necessary for up-regulation of glycolysis upon inhibition of oxphos.

Given that CK666 is added at the same time as CCCP, and inhibits both ADA and the initial ECAR increase by CCCP (both occurring within 4-min), it is likely to us that ADA is the relevant population of actin filaments responsible for the ECAR increase. However, Arp2/3 complex plays roles in many cellular processes, so a more specific link between ADA and the glycolytic increase is needed. We have previously shown that the initial step in CCCP-triggered Arp2/3 complex activation is a rise in cytoplasmic calcium, dependent upon the mitochondrial sodium-calcium exchanger NCLX (Fung et al., 2022). We asked whether the NCLX inhibitor CGP37157 would affect the CCCP-induced glycolytic response. When applied simultaneously with CCCP, CGP37157 lowers ECAR to a similar extent as CK666 (Figure S5 A). Oligomycin also potently increases ECAR (Pike Winer and Wu, 2014). We tested the effects of CK666 and CGP37157 on oligomycin-stimulated ECAR at 2 mM glucose. Similar to their effects with CCCP, both CK666 and CGP37157 inhibit the oligomycin-stimulated ECAR increase (Figure S5 B). These results suggest that ADA is the relevant Arp2/3 complex-dependent actin population that stimulates glycolysis, as opposed to another Arp2/3 complex-dependent process.

In contrast to its effects on ECAR, the effects of CK666 on oxygen consumption rate (OCR) are minimal for CCCP, antimycin A, and rotenone. As expected (Brand and Nicholls, 2011), CCCP increases OCR, while antimycin A and rotenone decrease OCR (Fig. S5, C-E). Simultaneous treatment with CK666 has no clear effect on OCR under any conditions (Fig. S5, C-E). These results show that CK666 affects the activation of glycolysis, rather than altering oxidative phosphorylation.

As a second method to assess glycolysis over a longer time period, we assayed lactate in the culture medium. At 2 mM glucose, lactate levels are significantly elevated by CCCP, antimycin A or rotenone treatment over a 5-hr time course, but simultaneous addition of CK666 suppresses this increase (Fig. S6 A). In contrast, the effect of CK666 at 25 mM glucose is comparatively mild (Fig. S6 B).

We also used the lactate assay to assess the effect of CK666 on glycolysis under hypoxic conditions (1% oxygen). At 2 mM glucose, CK666 inhibits lactate production 2.21-fold under hypoxic conditions (Fig. 3 G, 5-hr timepoint) but only 1.15-fold in normoxia (Fig. S6 C). At 25 mM glucose, lactate production is similar in the presence or absence of CK666 in normoxic or hypoxic conditions (Fig. S6, D and E). These results suggest that Arp2/3 complex-mediated actin polymerization is important for the up-regulation of glycolysis under hypoxic conditions.

Finally, we examined the effect of oligomycin on ATP levels and ECAR at both 25 mM and 2 mM glucose in MEFs. At 25 mM glucose, oligomycin treatment for 10 min causes a 10.16 ± 8.3 % increase in cytoplasmic ATP, which is brought back to baseline by CK666 addition (1.0 ± 1.1 %) (Fig. S7 A). At 2 mM glucose, oligomycin causes a 3.4 ± 5.4 % decrease in cytoplasmic ATP (Fig. S7 B). Even this small change in cytoplasmic ATP is sufficient to cause significant activation of AMP-dependent protein kinase (AMPK) (Fig. S7 C), which we have shown to be an initial step in ADA activation (Fung et al., 2022). In low glucose, CK666 addition to oligomycin causes further reduction of ATP level, to 19.7 ± 7.5 % (Fig. S7 B). These results suggest that glycolysis supplies the vast majority of ATP at either 25 or 2 mM glucose, but that Arp2/3 complex-mediated actin is required for optimal glycolysis under low glucose conditions when mitochondrial ATP synthesis is inhibited. The Seahorse assays suggest that the relevant Arp2/3 complex-mediated actin is ADA, based on its inhibition by both CK666 and CGP37157 (Fig. S5 B).

### ETC protein depletion causes mitochondrially-associated actin filaments and actin-dependent glycolytic activation

We tested whether longer-term reduction of mitochondrial oxphos induced an ADA-like response. One method for inducing chronic oxphos reduction is depletion of mitochondrial DNA (mtDNA), which in mammals contains genes encoding essential subunits of Complex I, III, IV, and V (Vafai and Mootha, 2012). Treatment with a low-dose of ethidium bromide (EtBr) causes mtDNA depletion (Fernández-Moreno et al., 2016). EtBr treatment of MEFs causes progressive mitochondrial depolarization over several days, with complete depolarization (comparable to CCCP) by 10 days (Fig. 4 A; and Fig. S8 A). During this time, mitochondria adopt a circular conformation (Fig. S8 B). ADA-like filaments arise around mitochondria by day 2, and are still present after 10 days (Fig. 4, B and C; and Fig. S8 B). Although this mitochondrially-associated actin is persistently present over multiple days (Fig. S8 B), it is inhibited within 5 min of CK666treatment (Fig. S8 C). This result suggests that the mitochondrially-associated actin filaments in these cells are dynamic, turning over with a half-life of less than 5 min.

**Figure 4.**
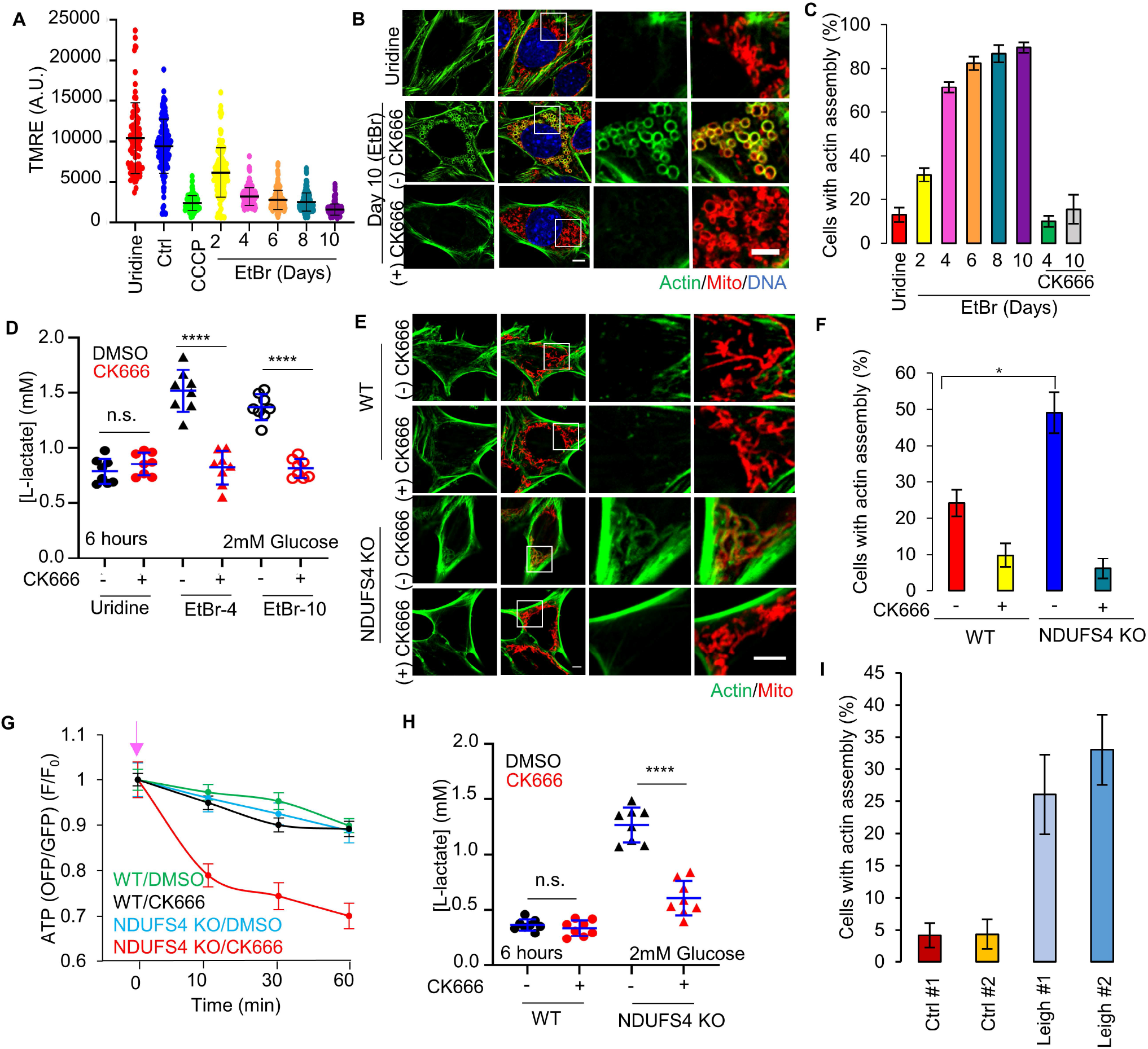
Actin assembly in ETC protein depleted cells. A) Mitochondrial polarization in MEFs (± s.d.) after 0.2 μg/ml ethidium bromide(EtBr)/50 μg/ml uridine treatment. Ctrl, untreated. Uridine, uridine treatment alone (10 days). CCCP - 10 min 20μM CCCP treated Ctrl cells. Circles indicate individual cell measurements (n ≥ 86 cells per group). Experiments done in 25mM glucose with serum. B) MEFs under uridine alone or ethidium bromide/uridine treatment (EtBr) for 10 days, stained for actin filaments (green) and mitochondria (red). Scale bar: 5 μm. C) % cells (± s.d.) displaying actin assembly after time in ethidium bromide/uridine, with and without 100μM CK666. n ≥ 98/15 cells/fields of view (FOV) per group. Experiments done in 25mM glucose with serum. D) Lactate production (± s.d.) in ethidium bromide cells (4 days, EtBr-4; 10 days, EtBr-10) and uridine-treated control (10 days), with and without 100μM CK666 after 6 hours. Points indicate individual well measurements starting with 75,000 cells/well. n.s. P > 0.05. **** P < 0.0001. Experiments done in 2mM glucose without serum. E) WT and NDUFS4 KO MEFs stained for actin filaments (green) and mitochondria (red). Scale bar: 5 μm. F) % cells (± s.d.) displaying actin assembly in WT and NDUFS4 KO MEFs, with and without 10 minutes of 100μM CK666 treatment. n ≥ 70/12 cells/FOVs per group. * P=0.018. Experiments done in 25mM glucose with serum. G) Cytosolic ATP levels in WT or NDUFS4 KO MEFs upon 100μM CK666 treatment. n ≥ 30 cells per group. Arrow indicates drug treatment. Experiments done in 25mM glucose with serum. H) Lactate production (± s.d.) in WT and NDUFS4 KO cells, with and without 100μM CK666 after 6 hours. n.s. P > 0.05. **** P < 0.0001. Points indicate individual well measurements starting with 75,000 cells/well. Experiments done in 2mM glucose without serum. I) % cells (± s.e.m.) displaying actin assembly in control or Leigh syndrome patient fibroblasts. n ≥ 76/13 cells/FOVs per group. Experiments done in 25mM glucose with serum. Statistical tests are tabled in Fig. S 10 B. Number of experiments, FOV, sample sizes and statistical tests are provided in Supplementary Table 1.

We examined the effect of these peri-mitochondrial actin filaments on glycolysis in EtBr-treated MEFs, testing lactate production in cells treated for either 4 and 10 days (EtBr-4 and EtBr-10 cells, respectively), and comparing to control cells treated with uridine only (control) for 10 days. In medium containing 2 mM glucose, lactate production is elevated in both EtBr-4 and EtBr-10 cells compared to control (Fig. 4 D; and Fig. S9, A-C). Treatment with CK666 reduces this lactate to control levels for both EtBr-4 and EtBr-10 (Fig. 4 D; and Fig. S9, A-C).

Another method for chronically reducing oxphos is knock-out of the NDUFS4 subunit of Complex I, which is associated with approximately 5% of autosomal recessive cases of the neurometabolic disorder Leigh syndrome (Lake et al., 2016; Rahman and Thorburn, 1993). Mice with NDUFS4 KO in neurons and glia display progressive encephalopathy that resembles the disease phenotype (Quintana et al., 2010). Examination of NDUFS4 KO MEFs reveals ADA-like peri-mitochondrial actin accumulation in the majority of cells (Fig. 4, E and F). Similar to mtDNA-depleted cells, this ADA-like actin is largely removed within 10 min of CK666 treatment (Fig. 4, E and F). These results suggest that longer-term inhibition of oxphos also leads to accumulation of actin around mitochondria.

We tested cytoplasmic ATP levels in NDUFS4 KO cells using the GO-ATeam1 sensor, suspecting that inhibition of ADA-like filaments would cause decreased ATP, similar to the mitochondrial poisons. In medium containing 2 mM glucose, treatment with CK666 causes an approximate 20% reduction in ATP levels in 10 min (Fig. 4 G), a similar time course to actin removal. In comparison, WT MEFs do not experience a significant ATP drop over 60 min of CK666 treatment (Fig. 4 G), similar to our earlier results. In terms of lactate production, NDUFS4 KO cells display characteristics similar to cells depleted of mitochondrial DNA. In medium containing 2 mM glucose, lactate production is significantly higher for these cells than WT MEFs, but is brought down to similar levels as WT MEFs by addition of CK666 (Fig. 4 H; and Fig. S9 D). In 25 mM glucose, CK666 treatment causes no significant change in lactate production for NDUFS4 cells (Fig. S9 E), again showing that the Arp2/3 complex-dependent effect on glycolysis does not occur under hyperglycemic conditions.

We also examined fibroblasts from Leigh syndrome patients for ADA-like actin accumulation around mitochondria. Cells from two patients with defined mutations were examined, in addition to cells from two control subjects. The two patient lines display a significant increase in the percentage of cells displaying peri-mitochondrial actin (Fig. 4 I; and Fig. S10), suggesting a similar situation to that in NDUFS4 KO cells.

These results suggest that, similar to the acute treatments, Arp2/3 complex-dependent actin polymerization is necessary for optimal glycolytic capability in cells that have chronic mitochondrial dysfunction. These cells also maintain polymerized actin around their mitochondria.

#### ADA-dependent glycolytic activation in effector CD8^+^ T cells

T cells undergo a dramatic metabolic change upon activation from naïve T cells to effector T cells (T_eff_), up-regulating glycolysis while also still using oxidative phosphorylation for significant ATP production (Geltink et al., 2018; Reina-Campos et al., 2021; Sena et al., 2013; van der Windt et al., 2012). Glycolytic activation is important for T_eff_ proliferation and the elaboration of effector functions to kill target cells(Chang et al., 2013; Menk et al., 2018). To test the importance of ADA in T cells, we isolated CD8^+^ T cells from the spleens of naïve mice, and activated them to T_eff_ *in vitro* with anti-CD3 and anti-CD28 antibodies. Treatment with CCCP, antimycin A, rotenone or hypoxia causes mitochondrially-associated actin polymerization in the majority of T_eff_, in a manner that is inhibited by CK666 (Fig. 5, A and B; and Fig. S11 A).

**Figure 5.**
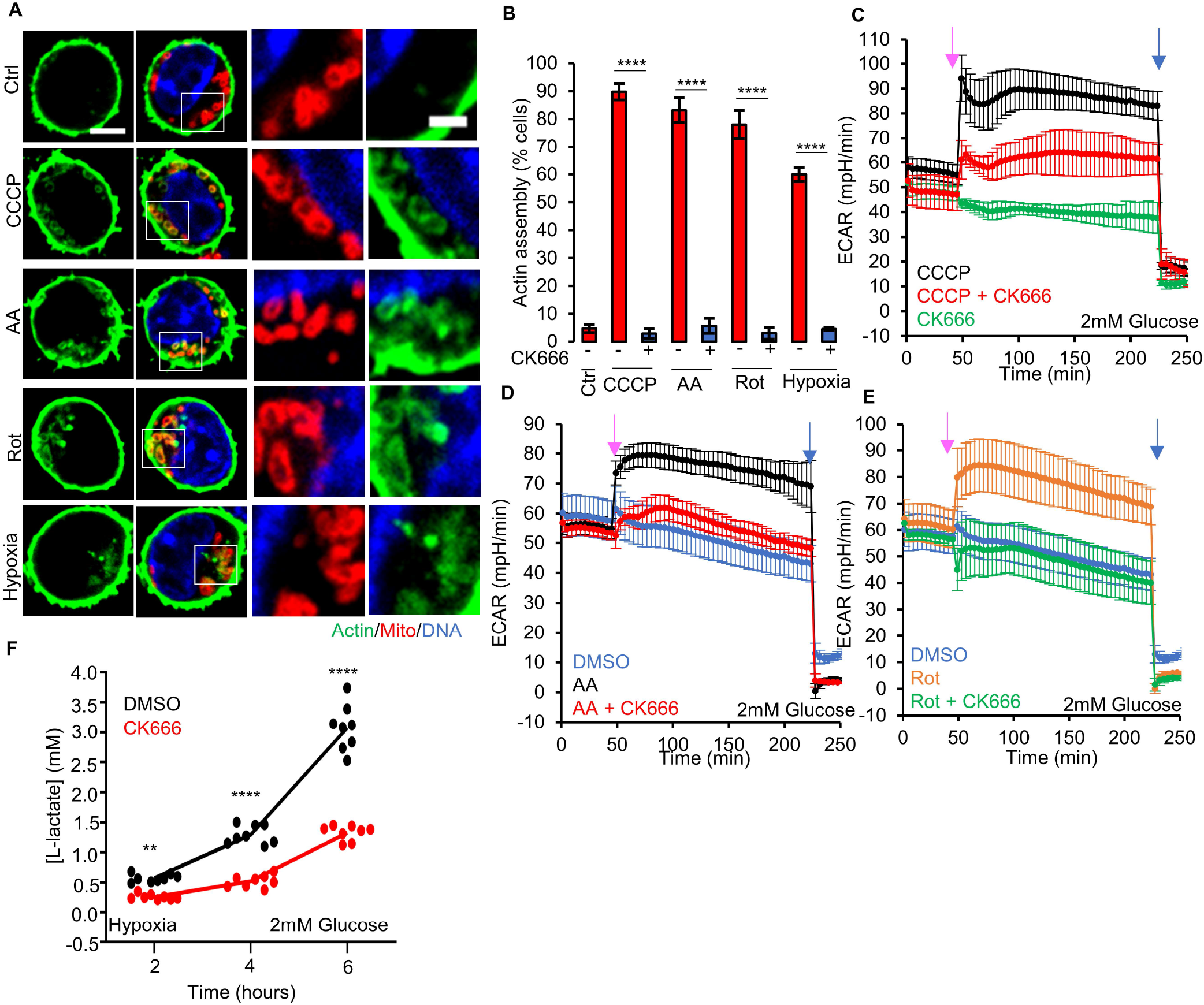
Effector T lymphocytes (T_eff_) require ADA for glycolytic activation. A) T_eff_ stained for actin filaments (TRITC-phalloidin, green), mitochondria (Tom20, red) and DNA (DAPI, blue) under un-stimulated conditions, or after treatment with 3 min 1μM CCCP, 5min 2.5μM antimycin A, 5 5μM min rotenone, or 60 min hypoxia in 2mM glucose medium. Right images are zooms of boxed regions. Scale bars: 5μm (full cell) and 2μm (inset). B) % cells (±s.e.m.) displaying ADA in treatments described in A, n ≥ 61/5 cells/FOV per group. **** P<0.0001. Experiments done in 2mM glucose without serum. C) ECAR (±s.d.) in T_eff_ upon addition of 100μM CK666, 1μM CCCP or 1μM CCCP + 100μM CK666 (45 min), followed by 50mM 2-deoxyglucose (2-DG)(223 min) in 2mM glucose medium without serum. Pink arrow indicates drug treatment and blue arrow indicate 2-DG treatment. D) ECAR (±s.d.) in T_eff_ upon addition of DMSO, 2.5μM antimycin A or 2.5μM antimycin A + 100μM CK666 (45 min), followed by 50mM 2-DG (223 min) in 2mM glucose medium without serum. Pink arrow indicates drug treatment and blue arrow indicate 2-DG treatment. E) ECAR (±s.d.) in T_eff_ upon addition of DMSO, 5μM rotenone or 5μM rotenone + 100μM CK666 (45 min), followed by 50mM 2-DG (223 min) in 2mM glucose medium without serum. Pink arrow indicates drug treatment and blue arrow indicate 2-DG treatment. F) Lactate production (2mM glucose without serum) induced by hypoxia (1% oxygen) in T_eff_ in the presence or absence of 100μM CK666 addition. Circles indicate individual well measurements starting with 400,000 cells/well. ** P=0.0013. **** P<0.0001. Number of experiments, FOV, sample size and statistical tests used are provided in Supplementary Table 1.

We then tested the effect of CK666 on glycolysis in T_eff_, using ECAR as a readout. At both 2 mM and 25 mM glucose, ECAR is stimulated by CCCP, antimycin A, and rotenone (Fig. 5, C-E; and Fig. S11, B-D). Interestingly, the ECAR response to antimycin A or rotenone treatment is rapid in T_eff_, in contrast to the slow response in MEFs. At 2 mM glucose, CK666 significantly inhibits the ECAR increase stimulated by all three treatments (Fig. 5, C-E), while the effects on OCR are unchanged (Fig. S11, E-G). At 25 mM glucose, simultaneous CK666 treatment reduces this ECAR increase slightly in all cases (Fig. S11, B-D).

T cells often encounter a hypoxic environment in solid tumors, and can be out-competed by highly glycolytic cancer cells under these conditions (Chang et al., 2015; Ho et al., 2015). We therefore tested the effect of hypoxia on glycolysis in T_eff_, using lactate production as a readout. At 2, 5 and 25 mM glucose, CK666 inhibits lactate production 2.32-, 1.75-, and 1.33-fold, respectively, under hypoxic conditions (Fig. 5 F; and Fig. S12, A and B, 6-hr timepoints). These results show that Arp2/3 complex-mediated actin polymerization stimulates glycolysis in T_eff_ upon treatments that compromise mitochondrial ATP production, with the effect being more pronounced at lower glucose concentration.

## DISCUSSION

In this paper, we show that actin polymerization is rapidly stimulated around mitochondria in response to multiple treatments that compromise mitochondrial ATP synthesis, including mitochondrial uncoupling (CCCP), inhibition of the electron transport chain (antimycin A, rotenone), inhibition of ATP synthase (oligomycin), and hypoxia. We refer to this actin accumulation as ADA, with the filaments being tightly apposed to the mitochondrion. A similar morphology of mitochondria-associated actin occurs under more chronic treatments that reduce mitochondrial oxidative phosphorylation, including mitochondrial DNA depletion, knock-out of the NDUFS4 subunit of complex 1 of the electron transport chain, and cells from Leigh syndrome patients. In all cases, inhibition of a key actin polymerization factor needed for ADA, Arp2/3 complex, inhibits the compensatory increase in glycolytic rate that occurs upon inhibition of mitochondrial ATP production. These results suggest that glycolysis is activated by peri-mitochondrial actin polymerization in response to decreased mitochondrial ATP synthesis.

The mechanism by which mitochondrial dysfunction induces ADA is intriguing. In response to CCCP, actin accumulates within 1-min and peaks by 4-min in multiple cell types. In the more chronic forms of mitochondrial dysfunction (EtBr treatment to deplete mtDNA, NDUFS4 KO, Leigh Syndrome cells), the peri-mitochondrial filaments are eliminated within 5 minutes of Arp2/3 complex inhibition, suggesting that this is not a pool of stably polymerized actin but is constantly turning over. We have previously shown that CCCP-induced ADA requires two parallel signaling pathways, one induced by increased cytoplasmic calcium, which activates Arp2/3 complex; and the other through AMPK activation, which activates the FMNL family of formins (Fung et al., 2022). One possibility is that mitochondrial depolarization is the initiating stimulus of these events. However, we show here that ADA-inducing stimuli span a wide range in terms of effects on mitochondrial polarization, including oligomycin, which causes slight hyper-polarization. Our current data might suggest that a decrease in mitochondrial ATP production capacity is a key signal. Even those stimuli that cause low changes in overall cytoplasmic ATP levels, such as oligomycin, cause significant and rapid AMPK activation, which might suggest an ability to detect ATP locally around mitochondria.

Another question concerns whether the mitochondrially-associated actin filaments induced during ADA are the cause of the glycolytic increase. While Arp2/3 complex mediates many actin-dependent cellular processes (Chakrabarti et al., 2021; Gautreau et al., 2021), three items suggest that ADA specifically contributes to glycolytic activation. First, inhibition of the mitochondrial sodium-calcium antiporter NCLX by CGP37157 inhibits the glycolytic increase caused by either CCCP or oligomycin stimulation. We have previously shown that NCLX mediates an important initial step in the ADA activation pathway (Fung et al., 2022). Second, the effects of both NCLX or Arp2/3 complex inhibition (by CGP37157 or CK666) on glycolysis occur within 4-min, because simultaneous addition of these compounds with CCCP inhibit the CCCP-induced ECAR increase at the first time point measured. While effects on other processes such as lamellipodia or endocytosis on this time scale are certainly possible, it is more likely to us that the inhibition of ADA de novo is the relevant event. Third, there is a strong inhibitory effect of CK666 on glycolytic activation by mitochondrial inhibitors in T_eff_, which have a limited number of existing actin-based structures. However, a role for other Arp2/3 complex-dependent actin processes in the rapid glycolytic activation we observe here cannot be ruled out.

The target linking Arp2/3 complex-mediated actin polymerization to this glycolytic increase is unclear, but a number of links between glycolysis and actin have previously been made. One study showed that aldolase was inhibited by an interaction with actin, and that insulin stimulation caused actin depolymerization and aldolase activation (Hu et al., 2016). Intriguingly, this insulin-stimulated aldolase activation was inhibited by CK-666, suggesting a more complicated mechanism than simply actin depolymerization. Whether this insulin-mediated glycolytic activation is related to the effects we observe is unclear, considering the different time courses of the responses (minutes for the effects reported here versus hours for the insulin effect). Another glycolytic enzyme that might be regulated by ADA is phosphofructokinase (PFK), whose degradation has recently been shown to be regulated by the E3 ubiquitin ligase TRIM21, itself being activated upon release from stress fibers (Park et al., 2020). Again, it is not clear that the effects reported here follow this mechanism, both in terms of speed of response and the fact that stress fibers are fundamentally different from Arp2/3 complex-dependent structures (Blanchoin et al., 2014). In budding yeast, a number of glycolytic enzymes appear to bind and be activated by actin (Espinoza-Simón et al., 2020), but links with Arp2/3 complex-mediated actin have not been made.

The effect of ADA on glycolysis is particularly important at lower glucose concentrations. While normal blood glucose ranges from 4-6 mM, lower glucose concentrations are common in peripheral tissues. In particular, neuronal cells experience steady-state glucose levels of 2.4 mM, which rapidly drop to below 1 mM during ischemia (Silver and Erecińska, 1994). The tumor micro-environment also can experience extracellular glucose levels below 1 mM, and competition for glucose between cancer cells and tumoricidal T_eff_ compromises anti-tumor effects (Chang et al., 2015; Ho et al., 2015). Rapid cellular proliferation and poor vascular supply lead to hypoxia in tumors, compromising mitochondrial function. Induction of ADA in infiltrating T cells might be therefore crucial in maintaining T cell viability and anti-tumor immunity.

Glycolytic activation is one of at least two functions of ADA. We have previously shown that ADA inhibits the mitochondrial re-organization that occurs downstream of depolarization (Fung et al., 2022; Fung et al., 2019). This re-organization occurs within the first 30 min after depolarization, and involves a circularization of the mitochondrion, rather than mitochondrial division (De Vos et al., 2005; Minamikawa et al., 1999; Miyazono et al., 2018). We showed that circularization depends upon the inner mitochondrial membrane protease Oma1 (Fung et al., 2022; Fung et al., 2019), one of whose substrates is Opa1 (MacVicar and Langer, 2016). Inhibition of ADA enhances Opa1 processing as well as circularization (Fung et al., 2022; Fung et al., 2019), suggesting that ADA might be able to exert some form of regulatory control over Oma1. The purpose of these shape changes are unclear, but may be a prelude to mitophagy. In this respect, ADA might serve as a temporary brake on responses to mitochondrial damage, increasing glycolytic rate to maintain ATP levels and delaying the mitophagic response. It is not clear whether the mitochondrial circularization we observe upon EtBr treatment represents a similar process to the rapid circularization induced by mitochondrial depolarization.

The exact organization of the actin filaments induced during ADA is unclear, but the staining intensity suggests them to be bundles of filaments or tightly-packed networks. The fact that Arp2/3 complex is required for ADA would suggest that a dendritic network might be present (Gautreau et al., 2021). Similar Arp2/3 complex-dependent actin structures, termed actin ‘clouds’ have been observed around mitochondria in mitotic (Moore et al., 2021) and interphase cells (Moore et al., 2016) in the absence of treatment with mitochondrial-compromising drugs. These actin clouds cycle in waves around the cell, making a full rotation within 15 minutes. The ADA response, however, appears to differ from actin clouds, in that it is not inhibited by wiskostatin (Fung et al., 2022). In contrast, we have previously shown ADA to be dependent on the WAVE family of Arp2/3 complex activators (Fung et al., 2022), while WAVE1 knock down does not inhibit mitotic actin clouds (Moore et al., 2021). It is possible, though, that ADA and actin clouds have overlapping functions, and it would be interested to determine whether actin clouds are associated with increased glycolysis.

Another type of recently-identified mitochondrial actin structure is termed actin ‘tails’, which develop from actin clouds during mitosis and increase mitochondrial motility, to favor homogenous distribution of mitochondria between daughter cells (Moore et al., 2021). ADA differs from actin tails in that it does not extend to micron lengths from the mitochondrion. In addition, we have not observed an increase in mitochondrial motility during ADA. In fact, we have previously shown that ADA suppresses mitochondrial dynamics (Fung et al., 2019).

In addition to the rapid effects on actin and glycolysis upon treatment with mitochondrial inhibitors, cells that are chronically compromised for mitochondrial function also have peri-mitochondrial actin. Arp2/3 complex inhibition eliminates this peri-mitochondrial actin in 5-min in ethidium bromide-treated cells. In NDUFS4-KO cells, Arp2/3 complex inhibition eliminates peri-mitochondrial actin and causes a substantial ATP drop in 10-min (the first time point we tested). Thus, these cells appear to continuously polymerize actin around their compromised mitochondria, perhaps to continuously stimulate glycolysis.

Finally, ADA is not the only Arp2/3 complex-dependent process in which actin polymerizes around damaged mitochondria. A second phase of actin polymerization occurs 1-2 hrs post-damage, which appears to function in the mitophagic process (Hsieh and Yang, 2019; Kruppa et al., 2018). In addition, we have previously mentioned in this discussion the Arp2/3 complex-dependent mitochondrial actin polymerization identified around apparently un-damaged mitochondria at interphase (Moore et al., 2016) and during mitosis (Li et al., 2015; Moore et al., 2021), which possess a number of differences to ADA. Our conclusion at present is that multiple mechanisms for Arp2/3 complex-mediated actin polymerization around mitochondria exist, activated by distinct mechanisms for distinct purposes. One purpose of ADA is to promote rapid glycolytic up-regulation in the face of mitochondrial dysfunction, in order to maintain cellular ATP levels.

## Materials and Methods

### Cell culture

Wild-type human osteosarcoma U2-OS and human cervical cancer HeLa cells were procured from American Type Culture Collection (ATCC) and grown in DMEM (Corning, 10-013-CV) supplemented with 10% newborn calf serum (NCS) (Hyclone, SH30118.03) for U2-OS or 10% fetal bovine serum (FBS)(Sigma-Aldrich F4135) for HeLa. Primate *Cercopithecus aethiops* Cos-7 cells were procured from ATCC (CRL-1651) and grown in DMEM with 10% FBS. Wild-type mouse embryonic fibroblasts (MEFs) were a gift from David Chan (previously described(Losón et al., 2013), while NDUFS4 KO MEFs were a gift from Yasemin Sancak (University of Washington, Seattle). MEFs were grown in DMEM with 10% FBS. Cell lines are cultivated at 37°C with 5% CO_2_ and were tested every 3 months for mycoplasma contamination using Universal Mycoplasma detection kit (ATCC, 30-1012K) or MycoAlert Plus Mycoplasma Detection Ki (Lonza, LT07-701). Cell lines were used no more than 30 passages.

### Mice and CD8^+^ T cells

Female wild-type C57BL/6NCrl mice were obtained from Charles River Laboratories. Mouse CD8^+^ T cells were isolated from the mouse spleens using EasySep™ Mouse CD8^+^ T Cell Isolation Kit (STEMCELL Technologies, 19853A) according to manufacturer instructions. Isolated CD8^+^ T cells were stimulated for 2 days in 24 well culture plates coated overnight with 10μg/mL anti-CD3 antibody (clone 145-2C11, BioXCell) and 5μg/mL soluble anti-CD28 (clone 37.51, BioXCell) antibody with 25U/mL recombinant human IL-2 (National Cancer Institute) in the medium. RPMI 1640 with L-Glutamine (Corning, 10-040-CV) supplemented with 10% FBS (HyClone, SH30541.03), 10mM HEPES (Corning Cellgro, 25-060-Cl), 1x non-essential amino acids (Corning Cellgro, 25-025-Cl), 1mM sodium pyruvate (Corning Cellgro, 25-000-Cl), and 44μM 2-Mercaptoethanol (Fisher Scientific, 03446I-100) was used for T cell stimulation and culture until the cells were harvested for experiments.

### EtBr treatment of MEFs

This treatment followed published protocols showing mtDNA depletion (Fernández-Moreno et al., 2016). 2 x10^4^ MEFs were plated directly onto Mat-tek imaging dishes and incubated in DMEM + 10% FBS overnight. 24 hours later, overnight media was replaced either with EtBr-containing media (DMEM + 10% FBS + 0.2 μg/ml EtBr (VWR life science, X328) + 50 μg/ml uridine) or control media (DMEM + 10% FBS + 50 μg/ml uridine). At designated times, cells were stained with TMRE to record mitochondrial membrane potential, following which they were fixed and stained for actin (TRITC-phalloidin), mitochondria (Tom-20) and nuclear DNA (DAPI).

### Human control and Leigh syndrome fibroblasts

All the culture cell materials from study subjects were collected with informed consent of the parents or the patient, following the recommendation from the Helsinki University Hospital ethical review board. The control cell lines originate from subjects eventually deemed not to manifest a mitochondrial disease.

The fibroblast cultures, previously immortalized by retroviral transduction of E6/E7 proteins of human papilloma virus, were cultivated at 37°C with 5% CO_2_ in DMEM (Dulbecco’s Modified Eagle’s Medium, Lonza Cat. #BE12-614F) with 10% fetal bovine serum albumin (Gibco, Cat. #11550356), 1 X GlutaMAX Supplement (Gibco, Cat. #35050061) 50 mg/l uridine (Calbiochem Cat. #6680) and 50 U/ml penicillin/streptomycin antibody (Gibco, Cat. #15070063) with media change in three day intervals. Cells were passaged after reaching 80% confluency by washing with PBS (Dulbecco’s Phosphate Buffered Saline, Sigma-Aldrich, Cat. #D8537-6×500ML) and incubating in 1 X trypsin-EDTA (Gibco, Cat. #15400-054) in 37°C for 3 min prior to replating on two fresh 10 cm dishes. Cell lines were used no more than 30 passages.

### DNA transfections and plasmids

For plasmid transfections, cells were seeded at 4×10^5^ cells per well in a 35 mm dish at ∼16 hours before transfection. Transfections were performed in OPTI-MEM medium (Gibco, 31985062) using lipofectamine 2000 (Invitrogen, 11668) as per manufacturer’s protocol, followed by trypsinization and re-plating onto glass-bottomed dishes (MatTek Corporation, P35G-1.5-14-C) at ∼1×10^5^ cells per well. Cells were imaged ∼16–24 h after transfection. GFP-F-tractin plasmid were gifts from C. Waterman and A. Pasapera (National Institutes of Health, Bethesda, MD) and were on a GFP-N1 backbone (Clonetech), as described previously (Johnson and Schell, 2009). Mito-DsRed construct was previously described (Korobova et al., 2014) and consist of amino acids 1–22 of *S. cerevisiae* COX4 N terminal to the respective fusion protein. ATP FRET sensor GoATeam1 was a gift from Hiromi Imamura (Kyoto University) and is described elsewhere (Imamura et al., 2009). The following amounts of DNA were transfected per well (individually or combined for co-transfection): 500 ng for Mito–DsRed, GFP–F-tractin and GoATeam1.

### Immunofluorescence

For all cell types and conditions, cells were fixed with 1% glutaraldehyde (Electron Microscopy Sciences, 16020) for 10 mins and subsequently washed three times with sodium borohydride (Fisher Chemical, S678)(1mg/ml, 15 mins interval) and then permeabilized with 0.25% TritonX-100 for 10 min. After permeabilization, they were washed thrice again with PBS and incubated in blocking buffer (10% NCS in PBS) for 30 min. The cells were then incubated with anti-Tom20 (Abcam, ab78547 1:500) antibody prepared in 0.1% blocking solution for 90 min. Following PBS washes, the cells were incubated with secondary antibody against Tom20 (Alexa Fluor 488-coupled anti-rabbit; Invitrogen #A11037; 1:200) mixed with TRITC-phalloidin (Sigma, P1951 1:400), 1X DAPI (Sigma, D9542) and incubated for 60 min. The cells were then washed with PBS, resuspended in 2 ml PBS and imaged on the same or following day.

#### Drug treatment

For drug treatments with MEFs, cells were seeded onto Mat-tek dishes at 200,000 cells/well and incubated at 37°C incubator overnight. Cells were fixed after 3mins treatment with 20μM CCCP (Sigma, C2759), 20μM CCCP + 100μM CK666 (Sigma, SML0006) simultaneously, 25μM antimycin A (Sigma; A8674), 25μM antimycin A + 100μM CK666 simultaneously, 50μM rotenone (Sigma; R8875), 50μM rotenone + 100μM CK666 simultaneously in serum-containing culture DMEM. For oligomycin treatments, cells were fixed after 5mins treatment with 1.5μM oligomycin (Sigma; 75351) or 1.5μM oligomycin + 100μM CK666 simultaneously in serum-free 2mM glucose DMEM. For EtBr and NDUFS4 KO MEFs, cells were treated with 100μM CK666 for 10 mins before fixation with glutaraldehyde. For human fibroblasts, cells were seeded onto μ-slide 8 well (ibidi, 80826) at 200,000 cells/well and incubated at 37°C incubator overnight. Cells were fixed without treatment to identify actin structures around mitochondria.

For drug treatments with effector T cells (T_eff_), cells were seeded onto Mat-tek dishes at 200,000 cells/well in serum-free (low glucose) DMEM medium (Agilent Seahorse XF DMEM; 103575-100) supplemented with 4mM L-glutamine, 1mM sodium pyruvate with 2mM D-glucose and allowed to adhere for 1 hour. Cells were fixed after 3mins treatment with 1μM CCCP, 1μM CCCP + 100μM CK666 simultaneously; or 5mins treatment with 2.5μM antimycin A, 2.5μM antimycin A + 100μM CK666 simultaneously, 5μM rotenone, 5μM rotenone + 100μM CK666 simultaneously. A control for no treatment was also performed.

#### Hypoxia

For hypoxia treatments, WT-MEF cells were seeded onto Mat-tek dishes at 200,000 cells/well and incubated at 37°C incubator overnight. Serum free DMEM medium was placed in the hypoxia chamber (InvivO2; BAKER; 94% N_2_, 5% CO_2_, 1% O_2_ at 37°C) overnight for equilibration. The following day, the Mat-tek dishes were washed twice with pre-warmed serum free DMEM. 2 ml of pre-equilibrated serum free DMEM (with DMSO, or CK666) were added to respective plates and quickly placed in the hypoxia chamber. 30 min later, the cells were fixed with glutaraldehyde and washed three times with PBS in the chamber before being taken out on the bench for downstream processing.

For effector T cells in hypoxia, serum free (low glucose) DMEM (Gibco, A1443001) with 2 mM D-glucose, 4 mM L-glutamine, 1mM sodium pyruvate was placed in the hypoxia chamber for equilibration overnight. The following day, cells were seeded onto a poly-D-lysine coated 8 well chamber slide (Ibidi, 80821) at 500,000 cells/well in low glucose and allowed to adhere in normoxia condition at 37°C for 1 hour. Next, the slide was brought into the hypoxia chamber and the pre-existing medium was substituted for the hypoxic medium containing either 100μM CK666 or DMSO. 1 hour later, the cells were fixed with glutaraldehyde and washed thrice with PBS before they were taken out of the chamber for sodium borohydride washes and subsequent steps.

### Antibodies and Western blotting

Anti-actin (mouse; mab1501R; Millipore) used at 1:1,000. Anti-phospho-AMPKα (Thr172) (CST;#2535; clone 40H9; rabbit monoclonal) was used at 1:1000. Li-COR secondary antibodies used were: anti-rabbit IRDye 800CW (#926-32211; 1:15000; goat) and anti-mouse IRDye 680RD (#926-68070; 1:15000; goat). For probing protein levels and AMPK phosphorylation in cell extracts, cells from a 35mm dish were trypsinized, centrifuged at 300 g for 5 min and resuspended in 400μl of 1× DB (50mM Tris-HCl, pH 6.8, 2mM EDTA, 20% glycerol, 0.8% SDS, 0.02% 784 Bromophenol Blue, 1M NaCl, 4M urea). Proteins were then separated by 10% SDS-PAGE in a Bio-Rad mini-gel system (7×8.4cm) and transferred onto polyvinylidene fluoride membrane (EMD Millipore, IPFL00010). The membrane was blocked with TBS-T (20 mM Tris-HCl, pH 7.6, 136 mM NaCl, 0.1% Tween-20) containing 3% BSA (VWR Life Science, VWRV0332) for 1h, then incubated with primary antibody solution at 4°C overnight. After washing with TBS-T, the membrane was incubated with fluorescently tagged Li-COR antibody for 1h at 23°C. Signals were detected by Li-COR fluorescent imager.

### Microscopy

Both fixed and live sample dishes were imaged using the Dragonfly 302 spinning disk confocal (Andor Technology) on a Nikon Ti-E base and equipped with an iXon Ultra 888 EMCCD camera, a Zyla 4.2M pixels CMOS camera, and a Tokai Hit stage-top incubator set at 37°C. A solid-state 405 smart diode 100 mW laser, solid state 560 OPSL smart laser 50 mW laser, and solid state 637 OPSL smart laser 140 mW laser were used. Objectives used were the CFI Plan Apochromat Lambda 100X/1.45 NA oil (Nikon, MRD01905) for all drug treatment in live-cell assays or fixed-cell assays. Images were acquired using Fusion software (Andor Technology, version 2.0.0.15). For live-cell imaging, cells were imaged in their respective cell culture medium. DMEM with 10% NCS for U2-OS or DMEM with 10% FBS for HeLa, Cos-7 and MEF cells. Medium was pre-equilibrated at 37°C and 5% CO_2_ before use. For actin burst and TMRE quantifications, cells were imaged at a single confocal slice at the medial region, approximate 2 μm above the basal surface, to avoid stress fibers. For live-cell drug treatments, cells were treated with 20μM CCCP, 20μM CCCP + 100μM CK666 simultaneously for media containing serum; or 1μM CCCP, 1μM CCCP + 100μM CK666 simultaneously for media free of serum at the start of the fifth frame (∼1min, with time interval set at 15s) during imaging and continued for another 9 min. To observe TMRE changes in WT, uridine treated or EtBr MEFs, cells were loaded with 20 nM TMRE for 30 min in culture medium at 37°C and 5% CO_2_. After incubation, TMRE was washed off with fresh culture medium before imaging.

### Image analysis and quantification

All image analysis was performed on ImageJ Fiji (version 1.51n, National Institutes of Health).

#### Immunofluorescence analysis

Cells with actin clouds were scored by visual analysis for the presence or absence of actin assembly in a given field and expressed as a percentage of the total number of cells for that field. Taking all the imaging fields into consideration for each condition, a bar graph showing the average percentage of cells, along with its respective errors bar in standard deviation (s.d.) or standard errors of the mean (s.e.m.) was plotted with Microsoft Excel. For human patient fibroblast dataset, imaging wells were blinded by P.W. Pallijeff and imaged by R. Chakrabarti. Microscopy images from each well were subsequently combined and scrambled by R. Chakrabarti before given to TS. Fung for analysis. Final quantification was unscrambled by R. Chakrabarti and then decoded by P. W. Pallijeff.

#### Quantification from live-cell imaging

The ImageJ Fiji Time Series Analyzer (UCLA) plugin was used for analysis for live-cell imaging. Cells that shrunk during imaging or exhibited signs of phototoxicity such as blebbing or vacuolization were excluded from analysis (maximal amount 10% for any treatment).

##### ADA

Quantification methods for actin assembly after mitochondrial-damage were previously described^3^. For each cell, one ROI was chosen which encompasses the entire area of ADIA around mitochondria after drug addition. Fluorescence values for each time point (*F*) were normalized with the mean initial fluorescence before drug treatment (first four frames−*F*_0_) and plotted against time as *F*/*F*_0_. For DMSO control or cells that did not exhibit an actin burst, the ROI was selected as the bulk region of the cytoplasm containing mitochondria using the mito– DsRed channel.

##### TMRE

Mean TMRE fluorescence was calculated from the entire mitochondrial area for each individual cell. TMRE fluorescence values for each time point (*F*) were normalized with the mean initial fluorescence before drug treatment (first four frames−*F*_0_) and plotted against time as *F*/*F*_0_.

### Cytoplasmic ATP changes using GO-ATeam1 biosensor

Go-Ateam1 plasmid (kindly provided by Hiromi Imamura, Kyoto University) was transfected into MEF cells using Lipofectamine 2000. The cells were then seeded onto Mat-tek dishes and imaged by spinning disk confocal microscopy. To acquire GFP and FRET signals, live cells were excited using a 488 nm laser and signals collected using a 525-50 nm band pass filter (GFP) and 600-50 nm band pass filter (OFP/FRET). Cells were imaged at a medial focal plane (i.e. not in the basal region of the cell) for 1-min at 15 s intervals to establish baseline fluorescence, then perfused with indicated drugs and continuously imaged for another 9-min. Mean fluorescence for GFP and OFP(FRET) channels, as well as OFP/GFP ratio, for a particular ROI (square of 50 μm^2^) from each cell (near the nucleus) were calculated for all time points using ImageJ. Ratios of each time point after drug treatment (F) were normalized with the mean initial ratio (first 5 frames before drug treatment) (F_0_) and plotted against time as F/F_0_. For oligomycin treated samples, data points were normalized to DMSO control curve.

### L-Lactate assay from extracellular medium

An assay kit based on the NADH-coupled reduction of tetrazolium salt to formazan (BMR service, University of Buffalo, SUNY, A-108) was used to measure the amount of L-lactate in the extracellular medium.

#### Drug treatment

For MEFs (wild type, uridine treated, NDUFS4 KO or EtBr) given drug treatments, cells were seeded at 75,000 cells per well in a 96 well plate and allowed to adhere overnight in culture medium. The following day, overnight medium was removed and cells were washed and equilibrated for 1 hour with either 1) high glucose medium - DMEM (Gibco, A1443001) with 25 mM D-glucose, 4 mM L-glutamine, supplemented with no serum or 2) low glucose medium - DMEM (Gibco, A1443001) with 2 mM D-glucose, 4 mM L-glutamine, supplemented with no serum. Next, cells were treated with 1μM CCCP; 1μM CCCP + 100μM CK666; 2.5μM AA; 2.5μM AA + 100μM CK666; 5μM Rotenone; 5μM Rotenone + 100μM CK666 for wild type cells. 100μM CK666 versus volume equivalent DMSO was used for wild type, NDUFS4 KO, uridine treated and EtBr cells. Cell culture medium was withdrawn at various timepoints (1-6 hours) and deproteinized in PEG solution. Samples were assayed with the L-lactate kit according to manufacturer’s instruction and OD 492nm was measured using a microplate reader (TECAN infinite M1000). A standard curve ranging from 0 to 1mM L-lactate was plotted for each experiment.

#### Hypoxia

For hypoxia experiments, MEFs were plated at 100,000 cells per well in a 96 well plate and allowed to adhere overnight at 37°C and 5% CO_2_ normoxia conditions while an aliquot of high/low glucose medium was placed inside the 1% O_2_ hypoxia chamber for oxygen depletion overnight. The following day, cells were brought into the hypoxia chamber, their pre-existing medium removed and treated with 100μM CK666 or DMSO containing hypoxic high/low glucose medium. Samples were taken at various timepoints (1-6 hours), deproteinized and assayed for L-lactate concentration.

For T_eff_ in hypoxic conditions, 400,000 cells were seeded per well in a poly-D-lysine treated 96 well plate and allowed to adhere and equilibrate for 1 hour at 37°C and 5% CO_2_ normoxia in high/low glucose medium (with additional 1mM sodium pyruvate) before being brought inside the hypoxia chamber. Cells were then treated with either CK666 or DMSO containing hypoxic high/low glucose medium. An additional condition with 5mM D-glucose also performed for T_eff_.

### ATP assays from cell extracts

A luciferase-based assay was used (BMR service, University of Buffalo, SUNY, A-125). MEFs were plated at 1 × 10^6^ cells per well in 60mm cell culture dishes and incubated for two days in standard medium (Corning DMEM containing 25 mM glucose, 10% FBS), resulting in 5.75 × 10^6^ cells/dish for MEFs. On the day of extraction, cells were washed and incubated for 1 hour in Agilent Seahorse XF DMEM (103575-100, with 4mM L-glutamine, no sodium pyruvate or serum) supplemented with 25mM glucose (high glucose condition) or 1mM glucose (low glucose condition) before treatment with: DMSO, 1μM CCCP; 100μM CK666 + 1μM CCCP; or 100μM CK666 for 2 mins. Cells were lysed immediately with 10% TCA and washed 3 times with 1:1 ether pre-saturated in TE (10mM Tris-HCl and 1mM EDTA, pH 8) for sample deproteinization. Samples were diluted 8-fold in water and ATP assay then assays conducted followed manufacturer’s instructions. The luminescence intensity was measured in a microplate reader (BioTek Synergy Neo2). A standard curve with ATP standards ranging from 0 -10μM was plotted for every experiment. The μM value of ATP determined in each assay was converted to a mM cellular value using the cell number stated above and an estimated cellular volume of 6 pL, obtained for Cos7 cells (Valm et al., 2017).

### Seahorse assay

A Seahorse XFe96 Bioanalyser (Agilent) was used to determine oxygen consumption rate (OCR) and extracellular acidification rate (ECAR) for MEFs and activated CD8^+^ cells. MEFs, taken from a T75 flask at 70-80% confluency and in culture for three days, were plated at 40,000 per well onto an Agilent XF microplate (101085-004) in DMEM (Corning; 10-013-CV) + 10% FBS medium, 24 hr before the experiment. Medium was changed 1-hour before the start of readout into 180 μL serum-free assay medium (Agilent Seahorse XF DMEM; 103575-100) supplemented with 4mM L-glutamine (Corning; 25-005-CI) and a variable amount of glucose (Agilent; 103577-100). The reading and injection regime was as follows: 1) Baseline OCR and ECAR were measured for 15min (6 measurements); 2) injection of CCCP (1μM final), CK666 (100μM final), or CK666 + CCCP; 3) measuring for 38 min (15 measurements); 4) injection of 50mM final 2-DG (Sigma Aldrich; D3179); 5) measuring for 18 minutes (7 measurements). Measurement parameters: 30 sec mix, 0 wait and 2 mins measure. For each experiment, three wells were done for CCCP or CK666 alone and four wells for CCCP + CK666. Stock concentrations: 10 mM CCCP (in DMSO), 20 mM CK666 (in DMSO), 500 mM 2-DG (in glucose free medium). CCCP and CK666 were diluted to 10x stocks in the appropriate medium and 20 mL injected (final DMSO concentration 0.5%). 22 mL 2-DG injected. “ΔECAR initial” represents the difference between the first measurement after the first injection and the last measurement before the injection, which represents a time span of approximately 2.5 min post-injection. “ΔECAR 40 min” represent the difference between the last measurement after the first injection and the last measurement before the first injection, which represents a time span of approximately 38 min post-injection.

For MEF treated with rotenone (Sigma; R8875) and antimycin A (Sigma; A8674), the reading and injection regime was as follows: 1) Baseline OCR and ECAR were measured for 30min (12 measurements); 2) injection of antimycin A (2.5μM final), CK666 (100μM final), or CK666 + antimycin A, rotenone (5μM final), or CK666 + rotenone or DMSO (volume equivalent to CK666); 3) measuring for 3 hours 22 mins (45 measurements); 4) injection of 50mM final 2-DG; 5) measuring for 30 minutes (12 measurements). Measurement parameters: 30 sec mix, 0 wait and 2 mins measure for baseline and 2-DG treatments, 30 sec mix, 0 wait and 4 mins measure for drug treatment. For each experiment, at least three wells were done for each condition. Stock concentrations for antimycin A and rotenone: 10 mM (in ethanol) and 10mM (in DMSO) respectively.

For MEF treated with oligomycin (Sigma; 75351) and CGP-37157 (CGP, Sigma; C8874), the reading and injection regime was as follows: 1) Baseline OCR and ECAR were measured for 24min (6 measurements); 2) injection of oligomycin (1.5μM final), CK666 (100μM final), or CK666 + oligomycin, CGP (80μM final), CGP + oligomycin, CCCP (1μM final), CK666 + CCCP, CGP + CCCP or DMSO (volume equivalent to oligomycin); 3) measuring for 1 hours 15 mins (15 measurements); 4) injection of 50mM final 2-DG; 5) measuring for 24 minutes (6 measurements). Measurement parameters: 2 min mix, 0 wait and 2 mins measure for baseline, drug and 2-DG treatments. For each experiment, at least three wells were done for each condition. Stock concentrations for oligomycin and CGP: 10mM (in DMSO) respectively.

For T_eff_, 150,000 cells (activated with anti-CD3 and anti-CD28 antibodies 48 hours prior) were plated into individual wells of a poly-D-lysine coated seahorse cell culture microplate (Agilent; 101085-004). T cell culture medium was changed 1 hour before the start of readout into 180μL serum-free assay medium (Agilent Seahorse XF DMEM; 103575-100) supplemented with 4mM L-glutamine, 1mM sodium pyruvate and either 2mM or 25mM glucose. The reading and injection regime was as follows: 1) Baseline OCR and ECAR were measured for 42min (12 measurements); 2) injection of various treatments, with the concentration same as MEFs: CCCP, CK666, CK666 + CCCP, antimycin A, CK666 + antimycin A, rotenone, CK666 + rotenone or DMSO (volume equivalent to CK666); 3) measuring for 2 hours 38 min (45 measurements); 4) injection of 50mM final 2-DG; 5) measuring for 42 minutes (12 measurements). Measurement parameters: 30 sec mix, 0 wait and 3 mins measure. For each experiment, at least three wells were done for each condition.

### Statistical analysis and graph plotting software

All statistical analyses and P value determinations were conducted using GraphPad Prism QuickCalcs or GraphPad Prism 9 (version 9.2.0, GraphPad Software). For P values in multiple comparisons (two way unpaired t-test; one or two-way ANOVA), Tukey’s multiple comparisons test was performed in GraphPad Prism 9. Scatter plots for TMRE, L-lactate assays were plotted with GraphPad Prism 9. ECAR and OCR curves, along with the standard deviation (s.d.); live-cell actin burst, along with the standard errors of the mean (s.e.m.) were all plotted using Microsoft Excel for Office 365 (version 16.0.11231.20164, Microsoft Corporation). Exact P values, sample size N and the number of independent experiments for each analysis are provided in Supplementary Table 1.

## Acknowledgements

We thank R. Cramer for use of the hypoxia chamber, T. Riggers for providing the stimulus, Z. Svindrych for his expertise in imaging, and N. Zaidi for help designing the Seahorse experiments. We thank J. Prudent, K. Rottner and Y. Sancak for sending us key reagents quickly. We thank Dr P. Isohanni for help collecting human patient cell samples. This work was funded by support from National Institutes of Health (NIH) grant R35 GM122545 (H.N. Higgs, R. Chakrabarti, TS. Fung), NIH grant P20 GM113132 (H.N.Higgs, R. Chakrabarti, TS. Fung) and NIH grant R01 AI155015 (E.J. Usherwood, T. Kang) and NIH grant R01 AI122854 (E.J. Usherwood, T. Kang).

## Author contributions

R. Chakrabarti, TS. Fung, E.J. Usherwood, A. Suomalainen and H.N. Higgs conceptualized the project and designed experiments. R. Chakrabarti, TS. Fung performed imaging experiments, T. Kang prepared and isolated effector T cells. P.W. Pallijeff prepared and isolated human patient fibroblasts. R. Chakrabarti, TS. Fung and T. Kang performed seahorse experiments. R. Chakrabarti and TS. Fung analyzed data. R. Chakrabarti, TS. Fung, P.W. Pallijeff, A. Suomalainen and H.N. Higgs designed the figures. E.J. Usherwood, A. Suomalainen, P.W. Pallijeff and T. Kang provided critical feedback. R. Chakrabarti, TS. Fung and H. N. Higgs wrote the paper and all authors contributed to methods and revisions.

## Correspondence and requests for materials

should be directed to H.N. Higgs.

## Supplementary Information

Supplementary Table 1 and Movies 1-5 are provided along with the manuscript.

## Figure Legends

**Figure S1.**
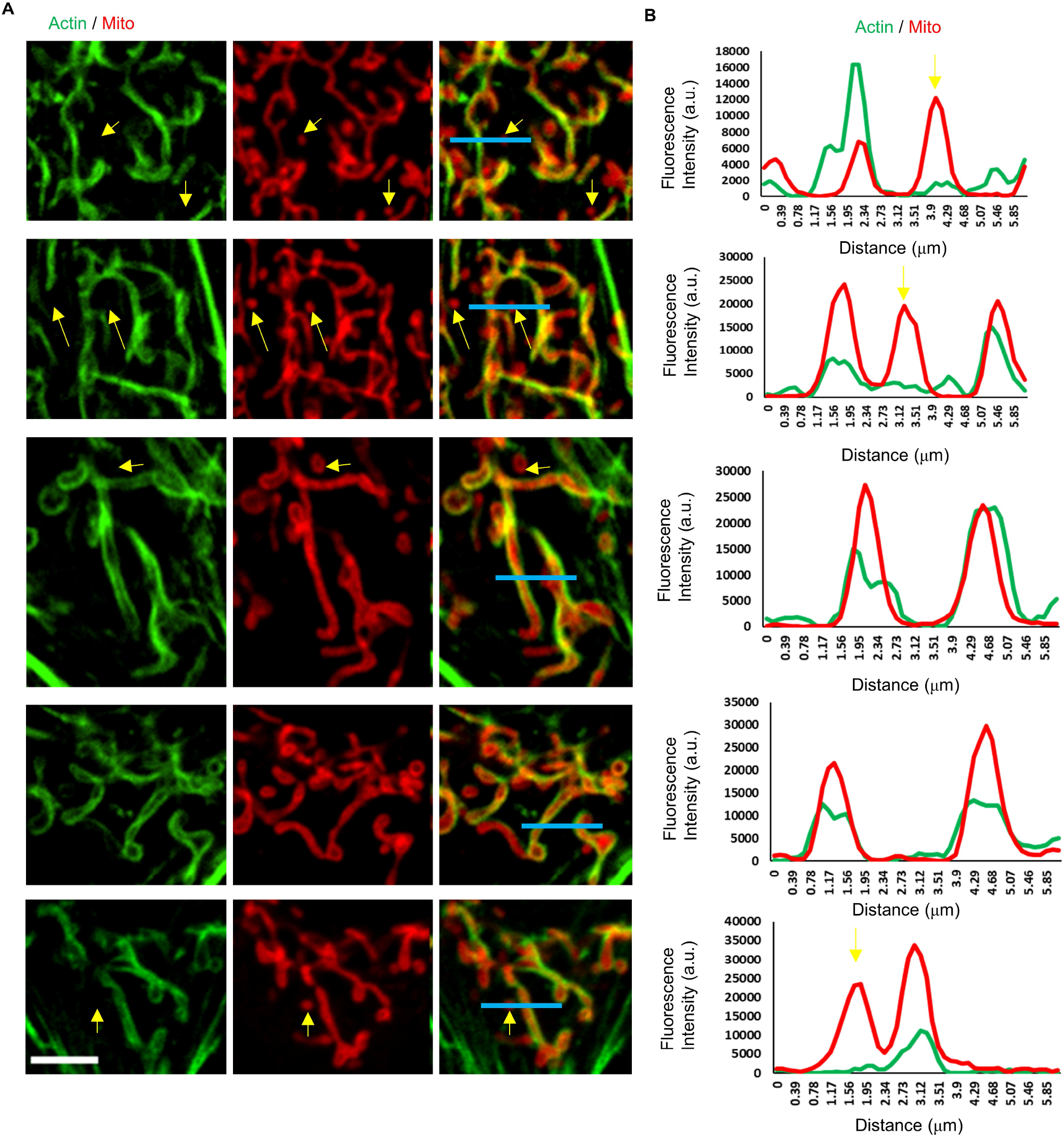
Line scans of ADA MEFs. A) Micrographs of actin assembly around mitochondria in fixed MEFs, stained for actin (green) and mitochondria (red). Scale bar: 5μm. Blue lines represent the region for line scans in B. Yellow arrows indicate punctate mitochondria without actin assembly. B) Line scans showing the fluorescent intensity for actin and mitochondria signal across each mitochondrion as shown in A.

**Figure S2.**
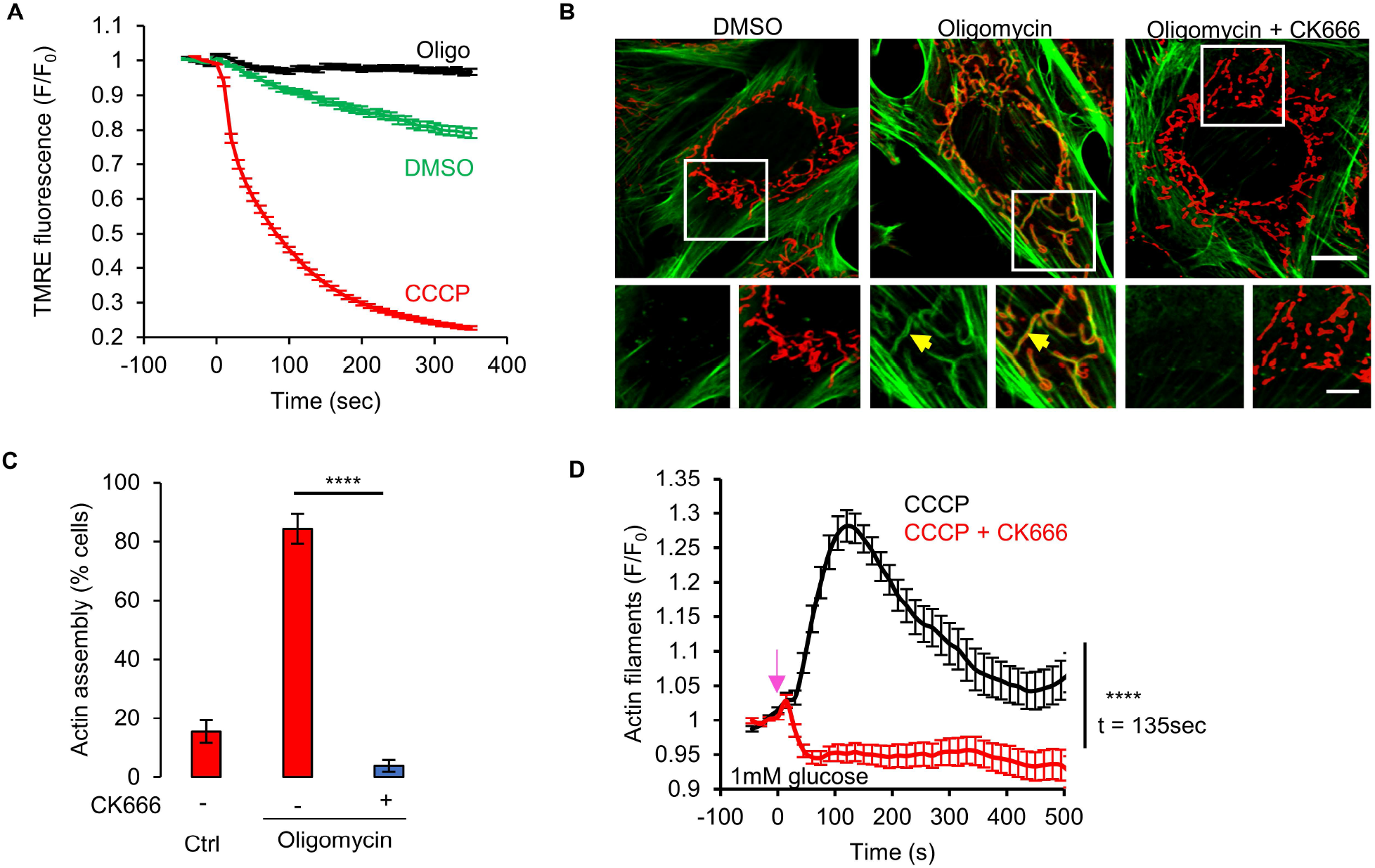
Oligomycin-induced ADA in MEFs. A) Mitochondrial polarization (assessed by TMRE fluorescence) in MEFs with DMSO, 1μM CCCP or 1.5μM oligomycin (± s.e.m.) treatment. n ≥ 118 cells per group. Experiments done in 2mM glucose without serum. B) MEFs stained for actin filaments (TRITC-phalloidin, green), mitochondria (Tom20, red) and DNA (DAPI, blue) after 5 min treatment with DMSO, 1.5μM oligomycin or 1.5μM oligomycin with 100μM CK666. Bottom images are zooms of boxed regions. Experiments done in 2mM glucose without serum. Scale bars: 10 μm and 5 μm. Arrow indicate actin assembly. C) % cells (± s.e.m.) displaying ADA for the conditions shown in panel B. n ≥ 65/14 cells/fields of view (FOV) per group. **** P<0.0001. Experiments done in 2mM glucose without serum. D) Graph of actin intensity (± s.e.m.) around mitochondria in MEF cells as a function of time for 1μM CCCP or 100μM CK666 + 1μM CCCP simultaneous treatment. Cells were cultured in Agilent seahorse DMEM supplemented with 1mM glucose and 4mM glutamine but without serum for 1 hour before imaging. n ≥ 35 cells per condition. Arrow indicates time of treatment. **** P<0.0001. Number of experiments, statistical tests and sample sizes are provided in Supplementary Table 1.

**Figure S3.**
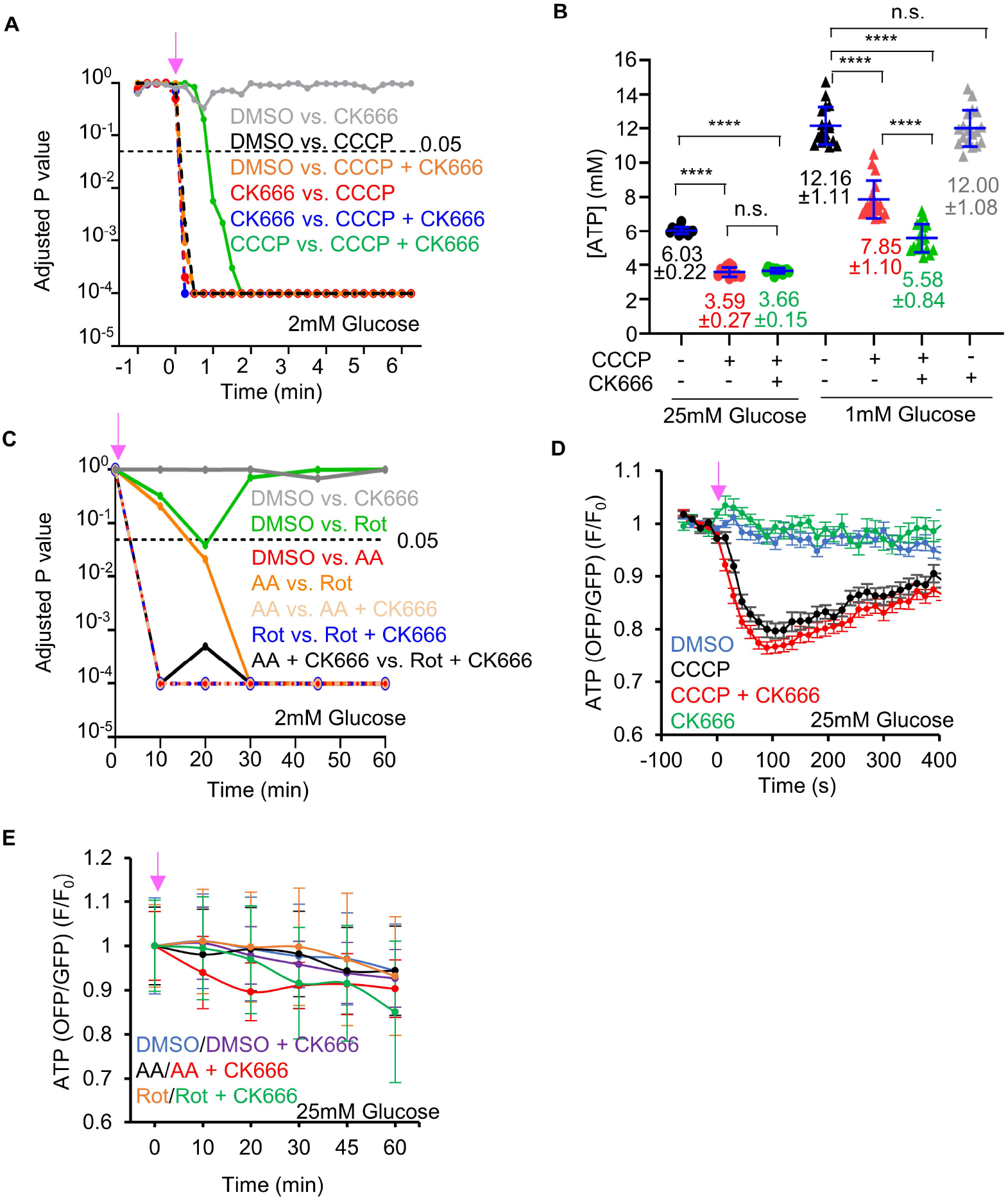
ATP levels changes by complex I or III inhibition. A) Graph of P values for comparisons of GO-ATeam1 timecourses in Fig 3 A (CCCP or CCCP/CK666 treatment of MEFs in 2mM glucose). B) ATP levels (± s.d.) in MEFs upon DMSO, 1μM CCCP or 1μM CCCP + 100μM CK666 treatments in medium containing either 1mM or 25mM glucose without serum, assayed from cell extracts. Points indicate individual measurements starting with 10^6^ cells/dish. N ≥ 12 measurements for each group. n.s. P > 0.05. **** P < 0.0001. C) Graph of P values for comparisons of GO-ATeam1 timecourses in Fig 3 B (antimycin A and rotenone treatments of MEFs in 2mM glucose). D) Graph of change in ATP levels (± s.e.m.) in live MEFs stimulated with 20μM CCCP in the absence or presence of 100μM CK666, using GO-ATeam1 biosensor. Cells cultured in medium containing 25mM glucose with serum. n ≥ 20 cells for each group. Arrow indicates time of treatment. E) Graph of change in ATP levels (± s.e.m.) in live MEFs stimulated with 25μM antimycin A or 50μM rotenone in the absence or presence of 100μM CK666, using GO-ATeam1 biosensor. Cells cultured in medium containing 25mM glucose with serum. n ≥ 33 cells for each group. Arrow indicates time of treatment. Number of experiments, statistical tests and sample sizes are provided in Supplementary Table 1.

**Figure S4.**
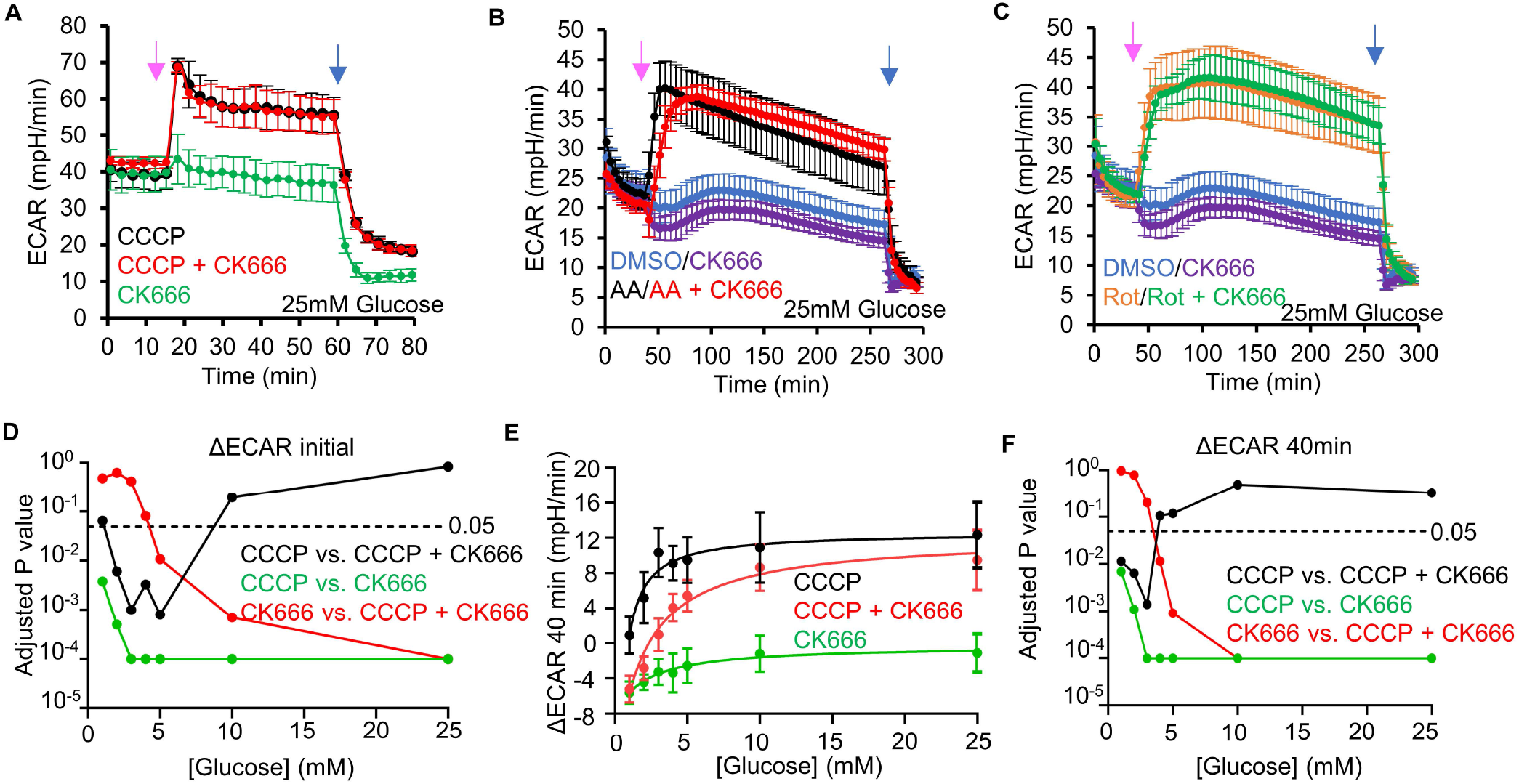
Changes in ECAR in MEFs after mitochondrial inhibitor treatments. A) ECAR (± s.d.) upon 100μM CK666, 1μM CCCP or 1μM CCCP + 100μM CK666 addition (15 min), followed by 50mM 2-DG (59 min) in 25mM glucose medium without serum. Pink arrow indicates drug treatment and blue arrow indicate 2-DG treatment. B) ECAR (± s.d.) upon DMSO, 100μM CK666, 2.5μM antimycin A or 2.5μM antimycin A + 100μM CK666 addition (33 min), then 50mM 2-DG (258 min) in 25mM glucose medium without serum. Pink arrow indicates drug treatment and blue arrow indicate 2-DG treatment. C) ECAR (± s.d.) upon DMSO, 100μM CK666, 5μM rotenone or 5μM rotenone + 100μM CK666 addition (33 min), then 50mM 2-DG (258 min) in 25mM glucose medium without serum. Pink arrow indicates drug treatment and blue arrow indicate 2-DG treatment. D) P values for comparisons between individual curves in Fig. 3 F. E) Effect of glucose concentration (± s.d.) on prolonged ECAR increase (after 40 min) induced by 1μM CCCP or 1μM CCCP + 100μM CK666 in MEFs. P values graphed in part F. F) P values for comparisons between individual curves in part E. Number of experiments, statistical tests and sample sizes are provided in Supplementary Table 1.

**Fig S5.**
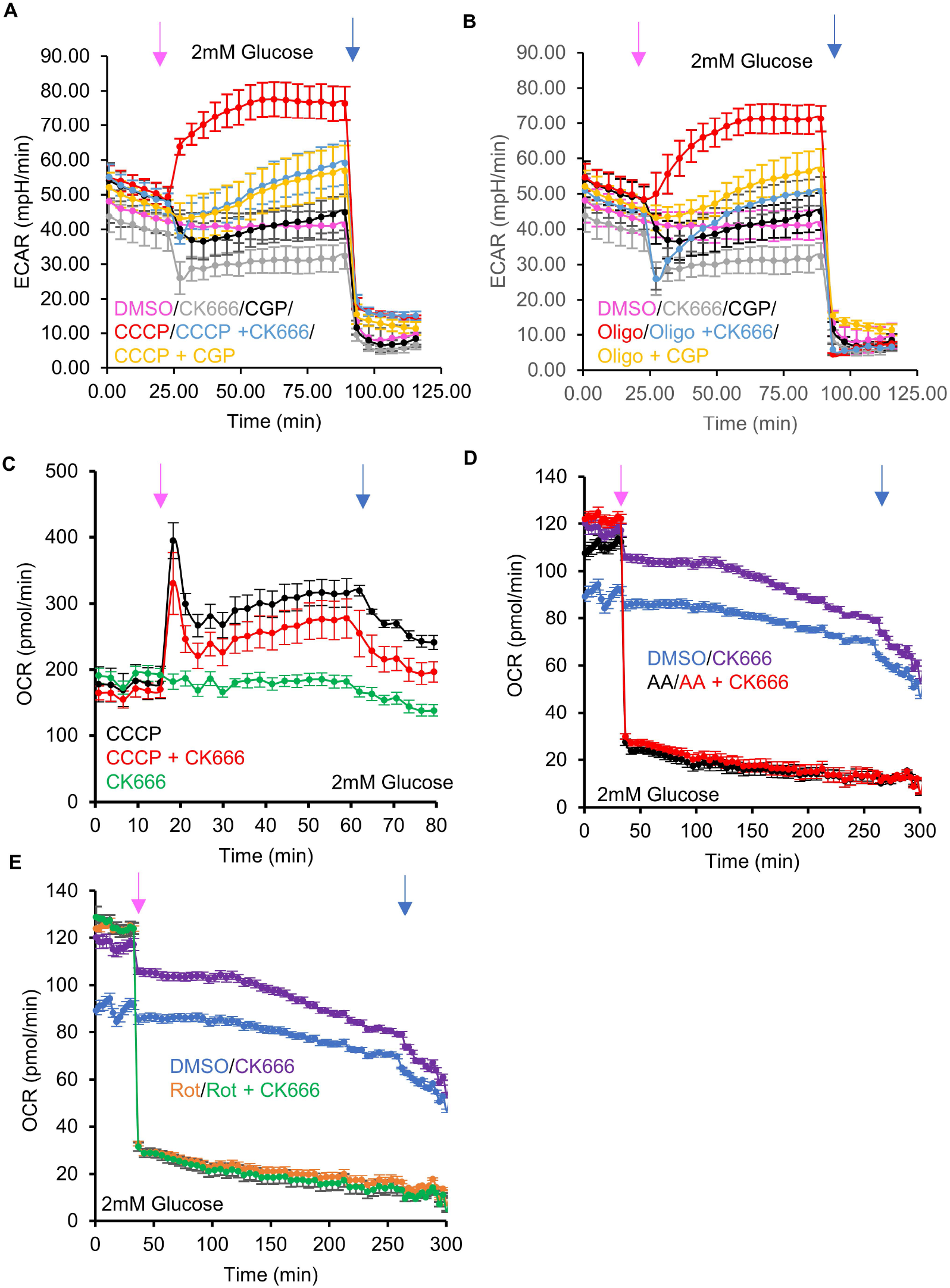
Effect of NCLX inhibition on CCCP- and oligomycin-activated glycolysis. A) ECAR (± s.d.) upon DMSO, 100μM CK666, 80 μM CGP37157, 1μM CCCP, 1μM CCCP + 100μM CK666 addition or 1μM CCCP + 80μM CGP37157 addition (at 23 min, pink arrow), followed by 50mM 2-DG (at 89 min, blue arrow) in 2mM glucose medium without serum. B) ECAR (± s.d.) upon DMSO, 100μM CK666, 80 μM CGP37157, 1.5μM oligomycin, 1μM oligomycin + 100μM CK666 addition or 1μM oligomycin + 80μM CGP37157 addition (at 23 min, pink arrow), followed by 50mM 2-DG (at 89 min, blue arrow) in 2mM glucose medium without serum. C) OCR (± s.d.) in MEFs (in 2mM glucose without serum) upon 100μM CK666, 1μM CCCP or 1μM CCCP + CK666 addition at 15 min, then 50mM 2-DG at 59 min. Pink arrow indicates drug treatment and blue arrow indicates 2-DG treatment. D) OCR (± s.d.) in MEFs (in 2mM glucose without serum) upon DMSO, 100μM CK666, 2.5μM antimycin A or 2.5μM antimycin A + 100μM CK666 addition at 33 min, then 50mM 2-DG at 258 min. Pink arrow indicates drug treatment and blue arrow indicates 2-DG treatment. E) OCR (± s.d.) in MEFs (in 2mM glucose without serum) upon DMSO, 100μM CK666, 2.5μM rotenone or 2.5μM rotenone + 100μM CK666 addition at 33 min, then 50mM 2-DG at 258 min. Pink arrow indicates drug treatment and blue arrow indicates 2-DG treatment. Number of experiments, statistical tests and sample sizes are provided in Supplementary Table 1.

**Figure S6.**
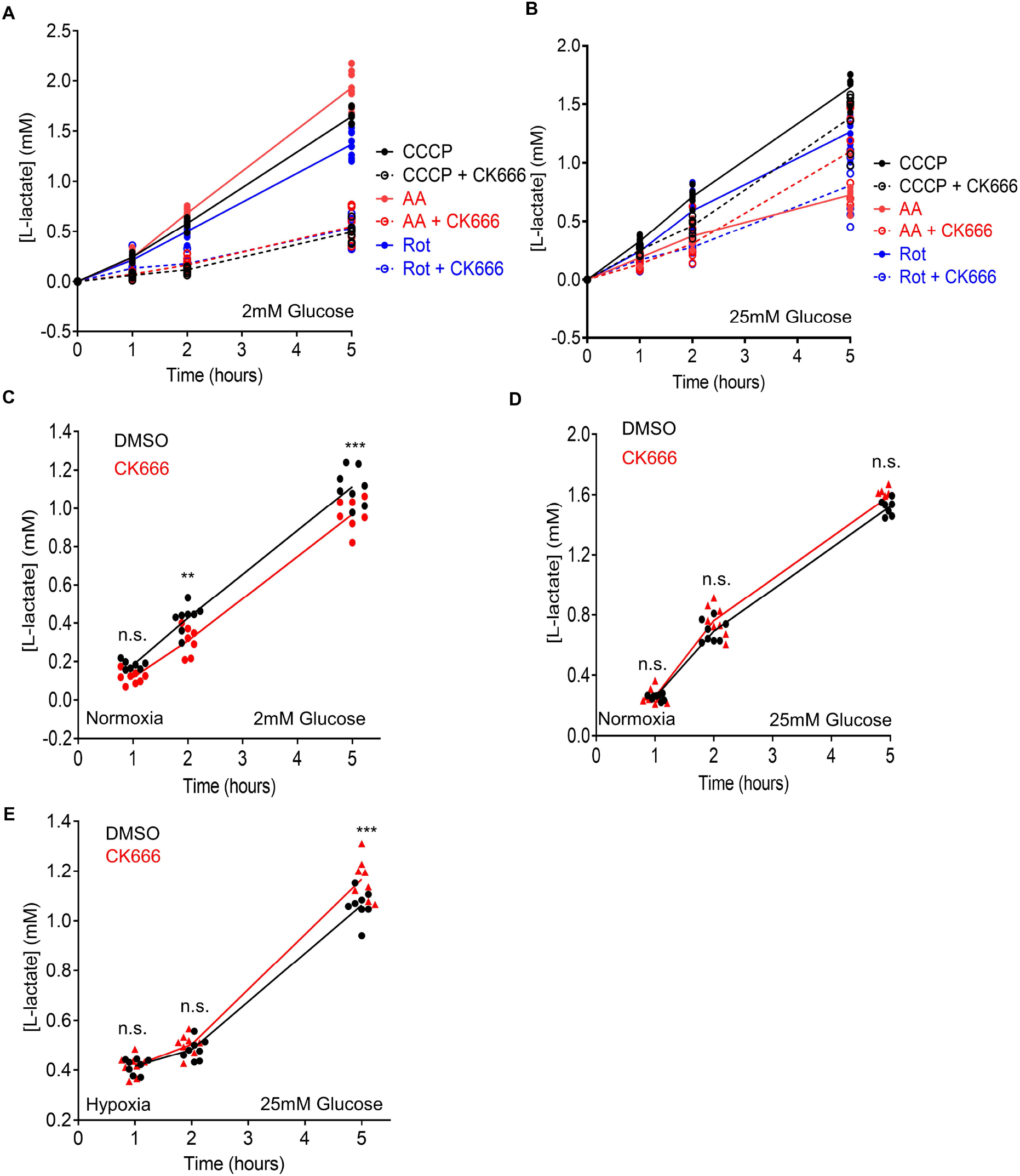
Changes in lactate production induced by mitochondrial inhibitors and hypoxia in MEFs. A) Effect of 100μM CK666 on lactate production upon 1μM CCCP, 2.5μM antimycin A, or 5μM rotenone treatment of MEFs in 2mM glucose without serum. Points indicate individual well measurements starting with 75,000 cells/well. B) Effect of 100μM CK666 on lactate production upon 1μM CCCP, 2.5μM antimycin A, or 5μM rotenone treatment of MEFs in 25mM glucose without serum. Points indicate individual well measurements starting with 75,000 cells/well. C) Effect of 100μM CK666 on lactate production in normoxia (21% O_2_) in MEFs at 2mM glucose without serum. Points indicate individual well measurements starting with 100,000 cells/well. n.s. P > 0.05. ** P = 0.002. *** P = 0.0002 D) Effect of 100μM CK666 on lactate production in normoxia in MEFs at 25mM glucose without serum. Points indicate individual well measurements starting with 100,000 cells/well. n.s. P > 0.05. E) Effect of 100μM CK666 on lactate production in hypoxia in MEFs at 25mM glucose without serum. Points indicate individual well measurements starting with 100,000 cells/well. n.s. P > 0.05. *** P = 0.0003 Number of experiments, statistical tests and sample sizes are provided in Supplementary Table 1.

**Figure S7.**
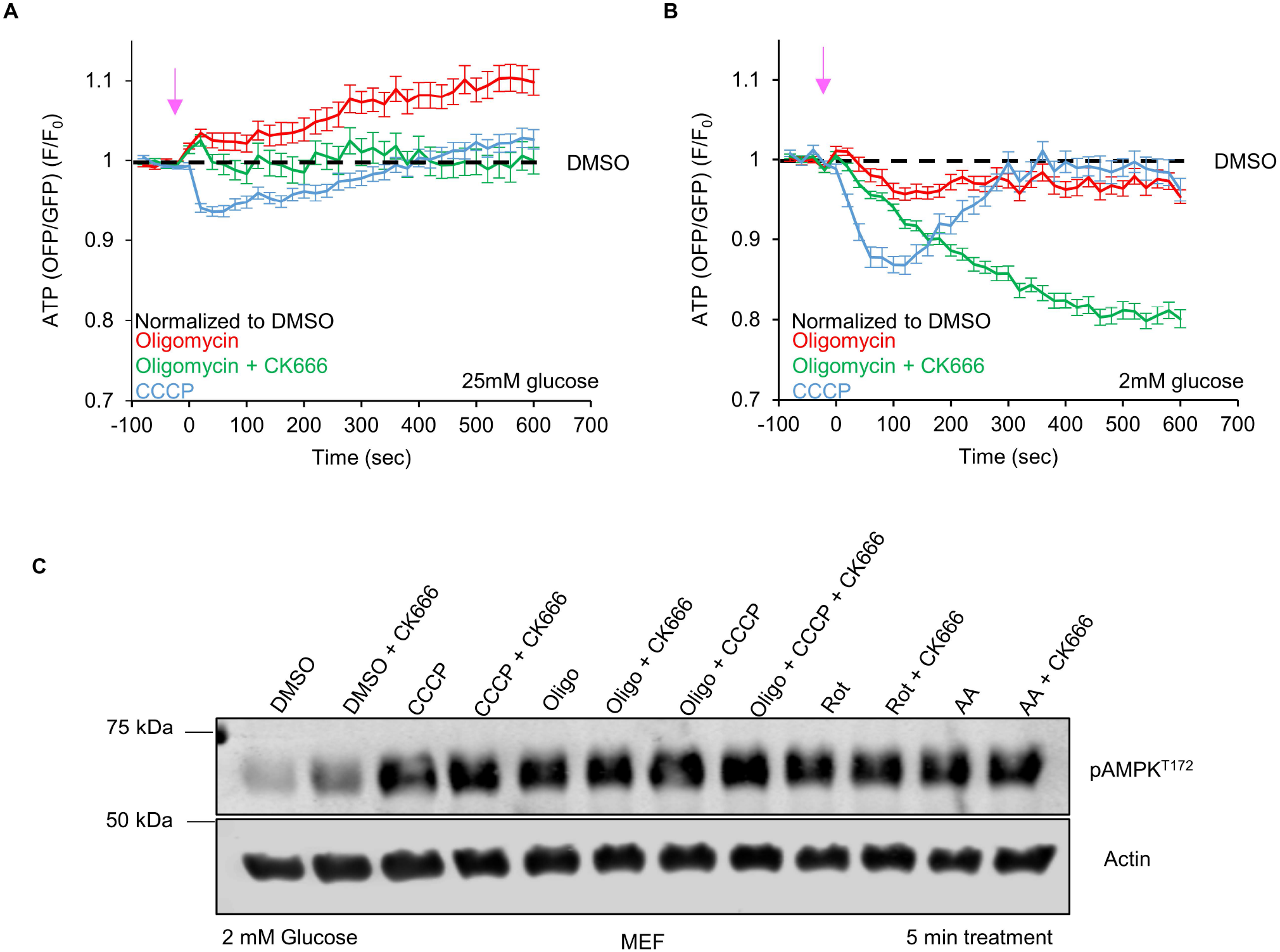
Cytoplasmic ATP changes induced by ATP synthase inhibition. A) Cytoplasmic ATP levels (± s.e.m.) after 1.5μM oligomycin in the absence or presence of 100μM CK666, or 1μM CCCP using GO-ATeam1. Data were normalized to DMSO control. n ≥ 24 cells per group. Arrow indicates time of treatment. Experiments done in 25mM glucose without serum. B) Cytoplasmic ATP levels (± s.e.m.) after 1.5μM oligomycin in the absence or presence of 100μM CK666, or 1μM CCCP using GO-ATeam1. Data were normalized to DMSO control. n ≥ 24 cells per group. Arrow indicates time of treatment. Experiments done in 2mM glucose without serum. C) AMPK activation after 5 min treatment of MEFs with ETC or ATP synthase inhibition. Actin is used as a loading control. Experiments done in 2mM glucose without serum. Number of experiments, statistical tests and sample sizes are provided in Supplementary Table 1.

**Figure S8.**
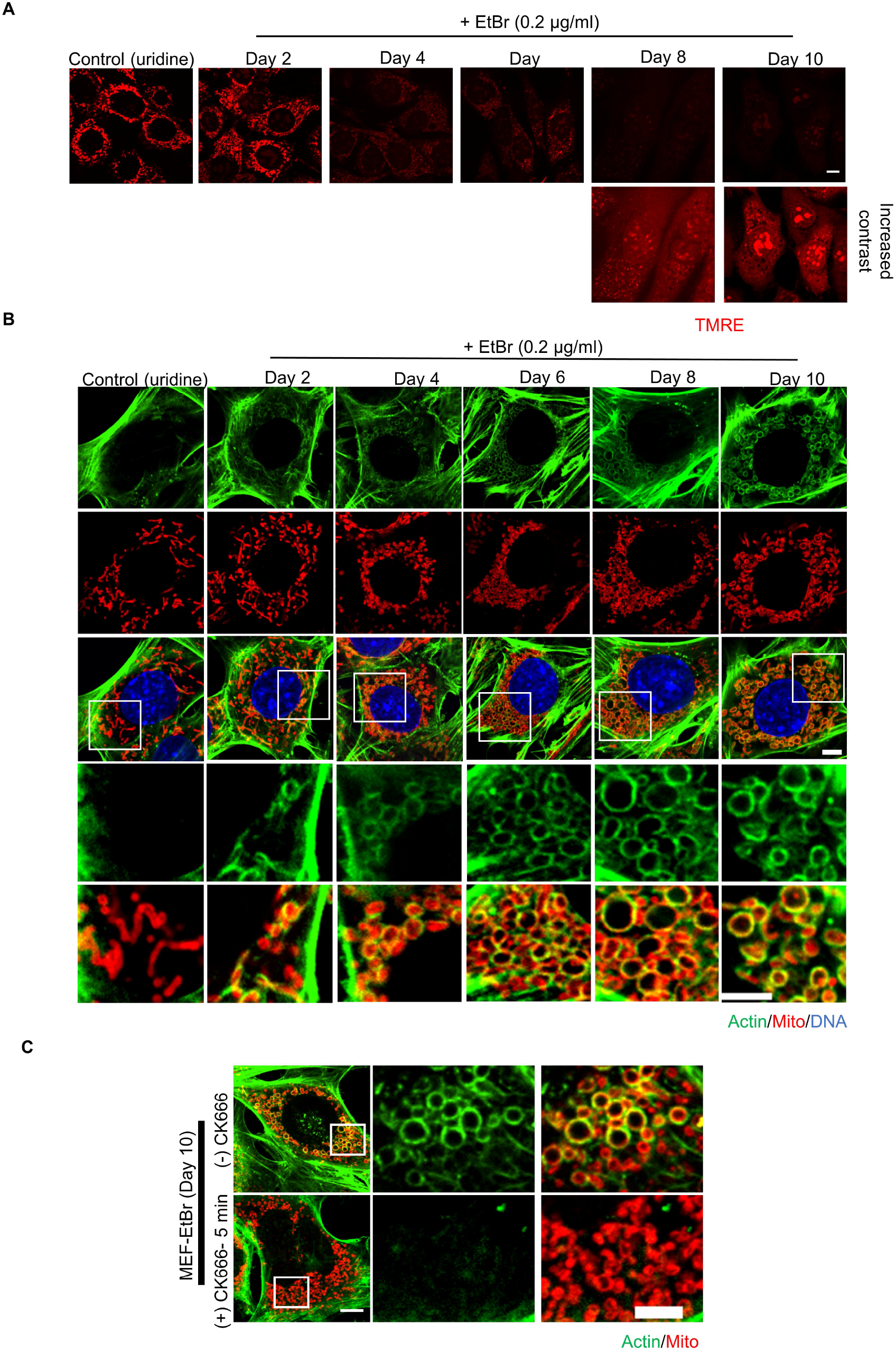
ADA and mitochondrial depolarization in EtBr-treated cells. A) Micrographs of TMRE staining of EtBr-treated cells at varying days post-treatment or control cells treated with uridine for 10 days. For days 8 and 10 post-EtBr treatment, the images below represent increased processing to reveal the presence of cells. Scale bar: 5 μm. B) Micrographs of actin staining (TRITC-phalloidin, green) around mitochondria (Tom20, red) at different days of EtBr treatment. DNA is stained with DAPI (blue). Images at the bottom are zooms of the boxed region. Scale bars: 5 μm. C) Micrographs of actin staining (TRITC-phalloidin, green) around mitochondria (Tom20, red) at day 10 of EtBr treatment with or without 100μM CK666 for 5min before fixation. Images at the right are zooms of the boxed region. Scale bars: 5 μm.

**Figure S9.**
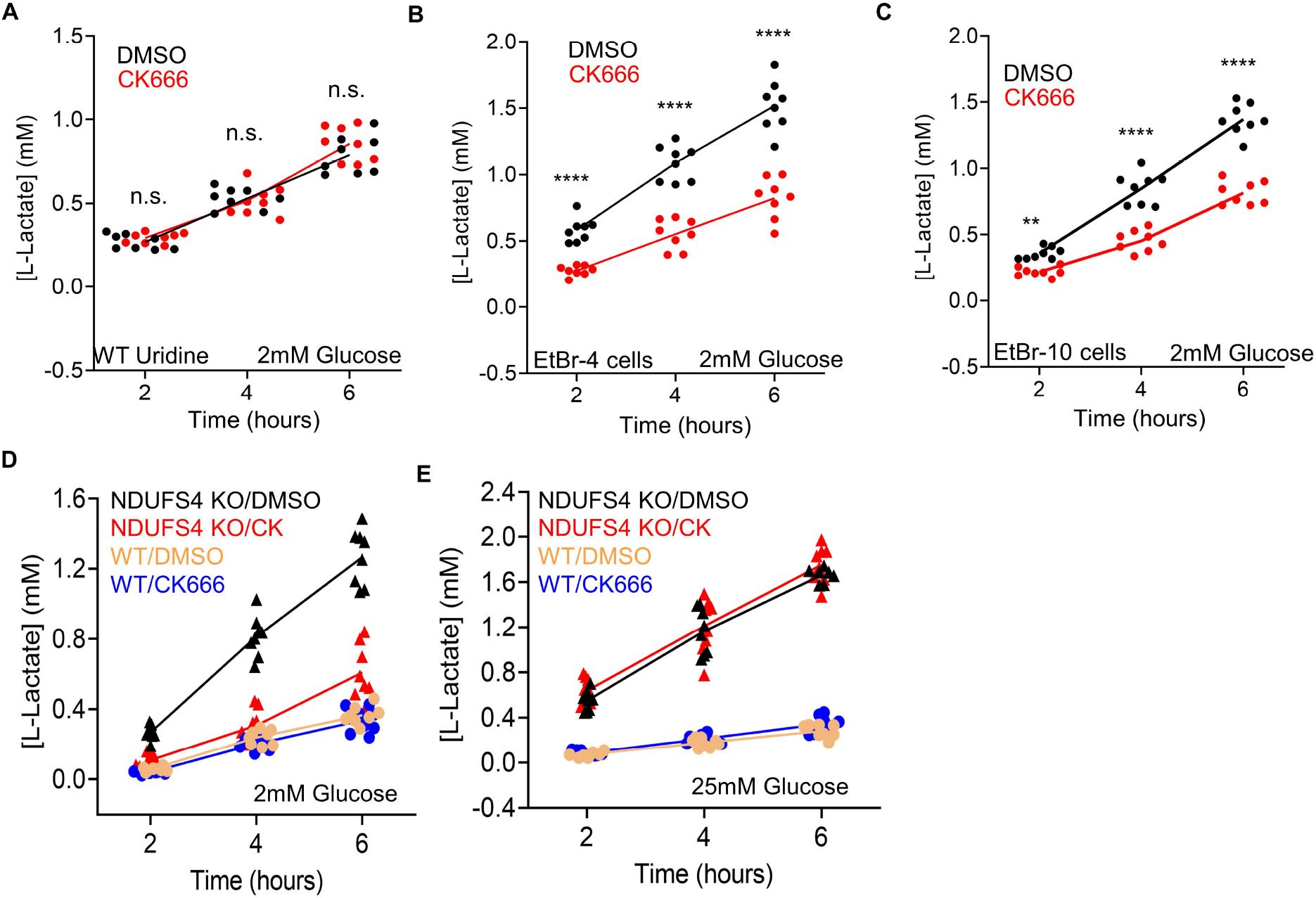
Changes in lactate production in EtBr or NDUFS4 KO MEFs. A) Time course of lactate production from control cells (uridine-treated and in 2mM glucose without serum) in the presence or absence of 100μM CK666. Points indicate individual well measurements starting with 75,000 cells/well. n.s. P > 0.05. B) Time course of lactate production from EtBr-4 cells (in 2mM glucose without serum) in the presence or absence of 100μM CK666. Points indicate individual well measurements starting with 75,000 cells/well. **** P < 0.0001. C) gTime course of lactate production from EtBr-10 cells (in 2mM glucose without serum) in the presence or absence of 100μM CK666. Points indicate individual well measurements starting with 75,000 cells/well. ** P = 0.0024. **** P < 0.0001. D) Time course of lactate production from WT and NDUFS4 KO cells (in 2mM glucose without serum) in the presence or absence of 100μM CK666. Points indicate individual well measurements starting with 75,000 cells/well. E) Time course of lactate production from WT and NDUFS4 KO cells (in 25mM glucose without serum) in the presence or absence of 100μM CK666. Points indicate individual well measurements starting with 75,000 cells/well. Number of experiments, statistical tests and sample sizes are provided in Supplementary Table 1.

**Figure S10.**
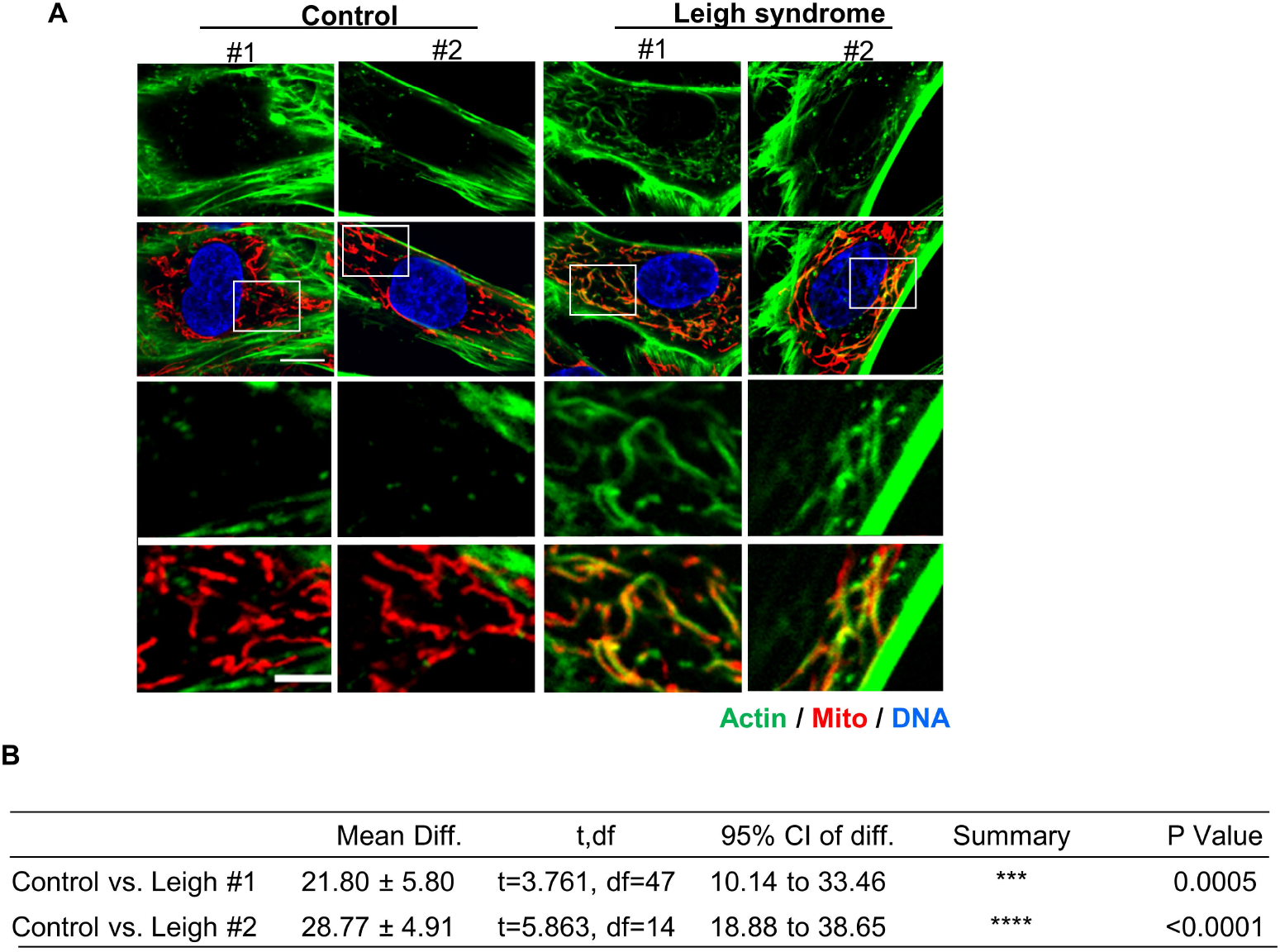
Actin assembly in Leigh syndrome fibroblasts. A) Micrographs of actin staining (TRITC-phalloidin, green) around mitochondria (Tom20, red) for control and Leigh syndrome fibroblasts. Scale bars are 10μm (full cell) and 5μm (inset). B) Table giving P values for comparisons of graph in Fig. 4I using unpaired student’s t test. Ctrl #1-2 were combined for analysis. Number of experiments and sample sizes are provided in Supplementary Table 1.

**Figure S11.**
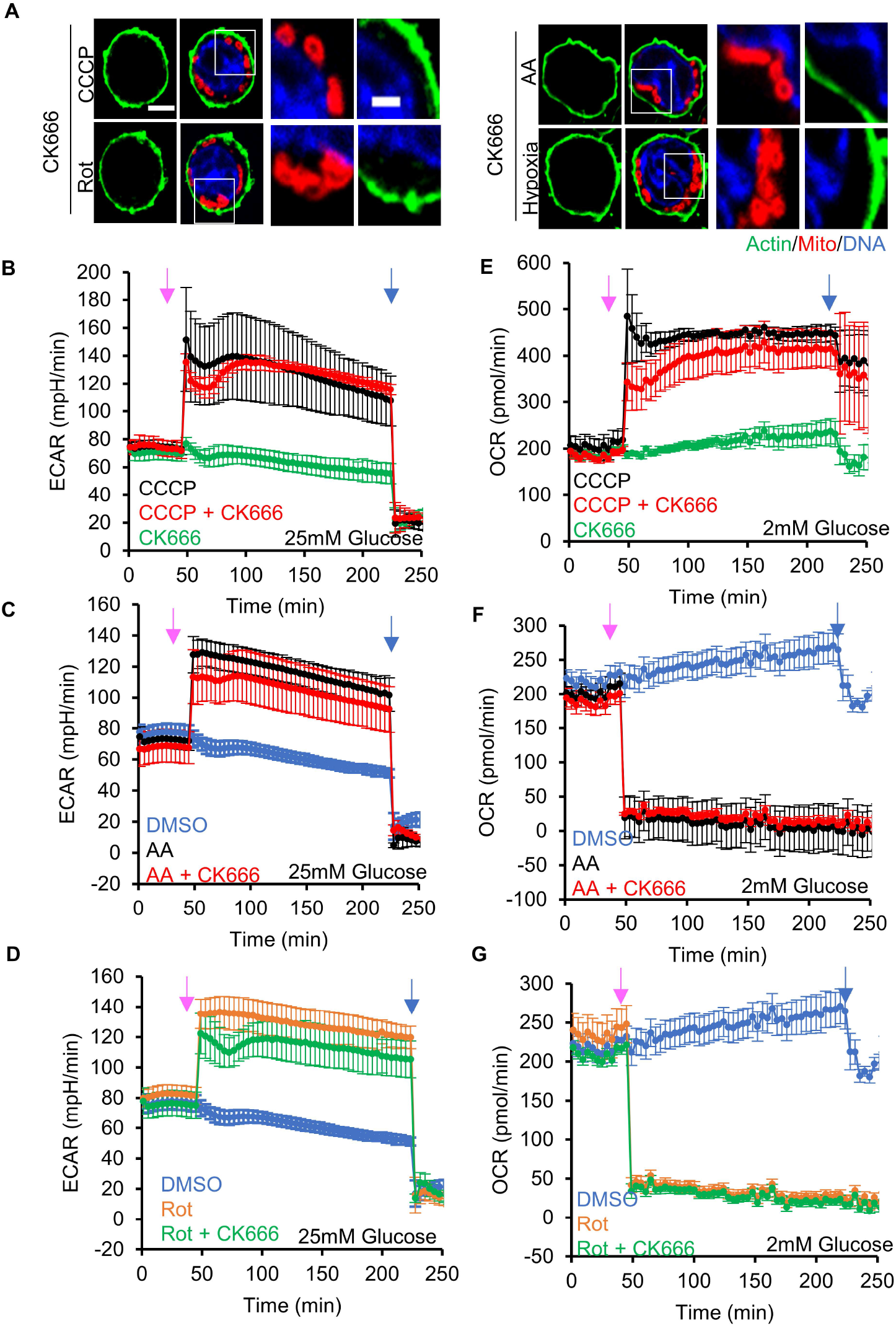
Effect of CK666 on ADA and glycolytic activation in T_eff_. A) T_eff_ stained for actin filaments (TRITC-phalloidin, green), mitochondria (Tom20, red) and DNA (DAPI, blue) after CCCP, antimycin A, rotenone, or hypoxia in the presence of CK666 for 2mM glucose medium without serum (1μM CCCP + 100μM CK666, 3 min; 2.5μM antimycin A + 100μM CK666 and 5μM rotenone + 100μM CK666, 5 min; hypoxia with 100μM CK666, 60 min). Images at right are zooms of boxed regions. Scale bars: 5μm (full cell) and 2μm (inset). B) ECAR (± s.d.) in 25mM glucose without serum upon 100μM CK666, 1μM CCCP or 1μM CCCP + 100μM CK666 addition at 45 min, then 50mM 2-deoxyglucose (2-DG) at 223 min. Pink arrow indicates drug treatment and blue arrow indicates 2-DG treatment. C) ECAR (± s.d.) in 25mM glucose without serum upon DMSO, 2.5μM antimycin A or 2.5μM antimycin A + 100μM CK666 addition at 45 min, then 50mM 2-DG at 223 min. Pink arrow indicates drug treatment and blue arrow indicates 2-DG treatment. D) ECAR (± s.d.) in 25 mM glucose without serum upon DMSO, 5μM rotenone or 5μM rotenone + 100μM CK666 addition at 45 min, then 50mM 2-DG at 223 min. Pink arrow indicates drug treatment and blue arrow indicates 2-DG treatment. E) OCR (± s.d.) in 2mM glucose without serum upon 100μM CK666, 1μM CCCP or 1μM CCCP + 100μM CK666 addition at 45 min, then 50mM 2-DG at 223 min. Pink arrow indicates drug treatment and blue arrow indicates 2-DG treatment. F) OCR (± s.d.) in 2mM glucose without serum upon DMSO, 2.5μM antimycin A or 2.5μM antimycin A + 100μM CK666 addition at 45 min, then 50mM 2-DG at 223 min. Pink arrow indicates drug treatment and blue arrow indicates 2-DG treatment. G) OCR (± s.d.) in 2 mM glucose without serum upon DMSO, 5μM rotenone or 5μM rotenone + 100μM CK666 addition at 45 min, then 50mM 2-DG at 223 min. Pink arrow indicates drug treatment and blue arrow indicates 2-DG treatment. Number of experiments and sample sizes are provided in Supplementary Table 1.

**Figure S12.**
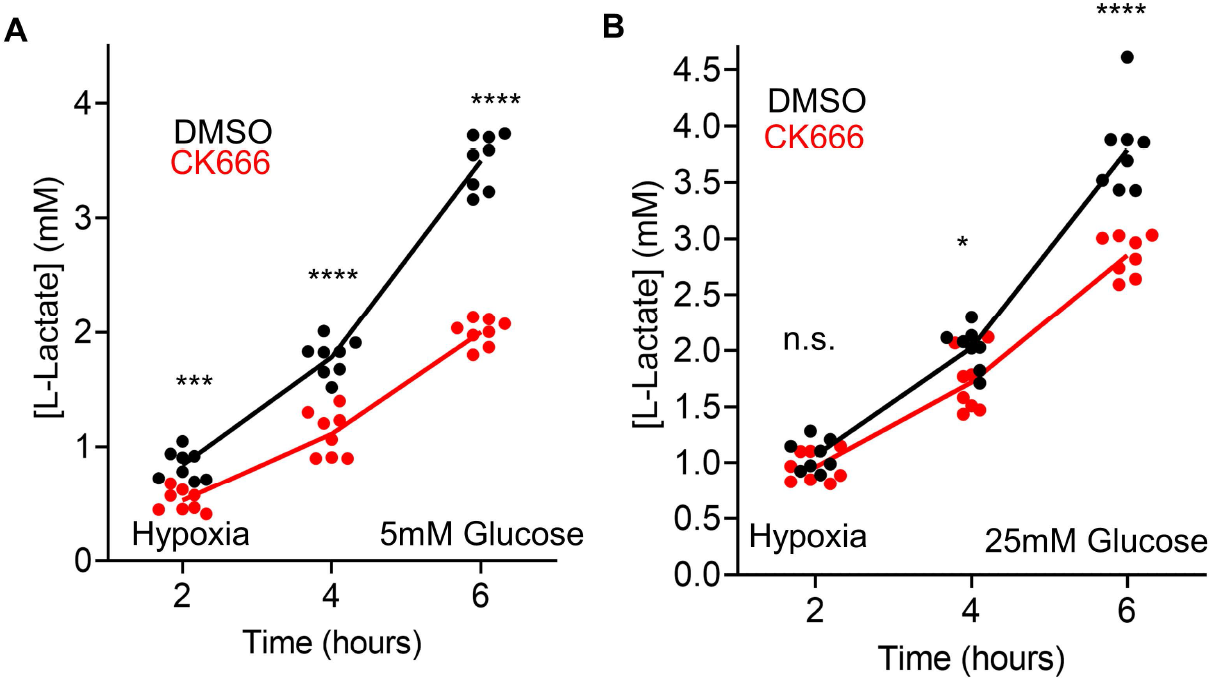
Effect of CK666 on hypoxia-induced lactate production in T_eff_. A) Lactate production induced by hypoxia (1% oxygen) in T_eff_ in the presence or absence of 100μM CK666 addition (5 mM glucose without serum). Circles indicate individual well measurements starting with 400,000 cells/well. *** P = 0.0003. **** P < 0.0001. B) Lactate production induced by hypoxia (1% oxygen) in T_eff_ in the presence or absence of 100μM CK666 addition (25mM glucose without serum). Circles indicate individual well measurements starting with 400,000 cells/well. n.s. P > 0.05. * P = 0.0136. **** P <0.0001. Number of experiments, statistical tests and sample sizes are provided in Supplementary Table 1.

## Movie legends

**Movie 1: ADA in U2-OS**.

U2-OS cell expressing GFP-F-tractin (green) and mito-BFP (red), stimulated with 20μM CCCP at time 00:00 (min:sec). Confocal images acquired every 15sec at medial region 2-4 μm above basal surface. Corresponds to Figure 1A. Scale bars: 10μm and 5μm (inset). Time stamp: min: sec.

**Movie 2: ADA in HeLa**.

HeLa cell expressing GFP-F-tractin (green) and mito-BFP (red), stimulated with 20μM CCCP at time 00:00 (min:sec). Confocal images acquired every 15sec at medial region 2-4 μm above basal surface. Corresponds to Figure 1A. Scale bars: 10μm and 5μm (inset). Time stamp: min: sec.

**Movie 3: ADA in Cos-7**.

Cos-7 cell expressing GFP-F-tractin (green) and mito-BFP (red), stimulated with 20μM CCCP at time 00:00 (min:sec). Confocal images acquired every 15sec at medial region 2-4 μm above basal surface. Corresponds to Figure 1A. Scale bars: 10μm and 5μm (inset). Time stamp: min: sec.

**Movie 4: ADA in MEF**.

MEF cell expressing GFP-F-tractin (green) and mito-BFP (red), stimulated with 20μM CCCP at time 00:00 (min:sec). Confocal images acquired every 15sec at medial region 2-4 μm above basal surface. Corresponds to Figure 1A. Scale bars: 10μm and 5μm (inset). Time stamp: min: sec.

**Movie 5: Zoomed-in view of ADA and mitochondria in U2-OS**

U2-OS cell expressing GFP-F-tractin (green) and mito-BFP (red) stimulated with 20μM CCCP at time 0 (sec). Confocal images acquired every 15sec at medial region 2-4 μm above basal surface. Scale bars: 2μm.Time stamp: sec.

